# Exploring the conformational changes of the Munc18-1/syntaxin 1a complex

**DOI:** 10.1101/2022.11.29.518383

**Authors:** Ioanna Stefani, Justyna Iwaszkiewicz, Dirk Fasshauer

## Abstract

Neurotransmitters are released from synaptic vesicles, the membrane of which fuses with the plasma membrane upon calcium influx. This membrane fusion reaction is driven by the formation of a tight complex comprising the plasma membrane SNARE proteins syntaxin-1a and SNAP-25 with the vesicle SNARE protein synaptobrevin. The neuronal protein Munc18-1 forms a stable complex with syntaxin-1a. Biochemically, syntaxin-1a cannot escape the tight grip of Munc18-1, so formation of the SNARE complex is inhibited. However, Munc18-1 is essential for the release of neurotransmitters *in vivo*. It has therefore been assumed that Munc18-1 makes the bound syntaxin-1a available for SNARE complex formation. Exactly how this occurs is still unclear, but it is assumed that structural rearrangements occur. Here, we used a series of mutations to specifically weaken the complex at different critical positions in order to induce these rearrangements biochemically. Our approach was guided through sequence and structural analysis and supported by molecular dynamics simulations. Subsequently, we created a homology model showing the complex in an altered conformation. This conformation presumably represents a more open arrangement of syntaxin-1a that permits the formation of SNARE complex to be initiated while still bound to Munc18-1. In the future, research should investigate how this central reaction for neuronal communication is controlled by other proteins.

## Introduction

Ca^2+^-dependent synaptic exocytosis is driven by the fusion of neurotransmitter-loaded synaptic vesicles with the presynaptic plasma membrane. The underlying membrane fusion reaction is catalyzed by the interaction of <u>N</u>-ethylmaleimide-<u>s</u>ensitive factor <u>a</u>ttachment <u>re</u>ceptors (SNAREs) with the plasma membrane, syntaxin-1a and SNAP25, and the synaptic vesicle, synaptobrevin (1). They zip into a tight four-helical bundle, known as the SNARE complex. SNARE proteins form a larger protein family that catalyzes the fusion of various types of transport vesicles in eukaryotic cells. Several other conserved proteins ensure that the SNARE proteins are tightly regulated, including the Rab proteins, <u>S</u>ec1/ <u>M</u>unc18-like (SM) proteins, and a group of tethering proteins known as <u>c</u>omplex <u>a</u>ssociated with tethering <u>c</u>ontaining <u>h</u>elical <u>r</u>ods (CATCHR) proteins (2-4).

The activity of the SNARE syntaxin-1a is tightly regulated by Munc18-1, an SM protein. Knock-out studies and mutational screens have shown that its interaction with syntaxin is indispensable for neurotransmission (5). *De novo* mutations in the Munc18-1 coding gene, STXBP1, have been associated with neurodevelopmental disorders (6, 7). *In vitro*, the strong interaction of Munc18-1 with syntaxin-1a inhibits the binding of the latter to its SNARE partners (8). A way to reconcile the biochemical inhibitory function with the positive genetic role has been sought for a long time. Many successes have been achieved, but, ultimately the final resolution of this apparent discrepancy is still pending (9-12).

The structure of the Munc18-1/syntaxin-1a complex (11) shows how Munc18-1 encloses and holds syntaxin-1a. Munc18-1 folds into an arch-shaped structure that wraps around syntaxin-1a in a so-called closed conformation, in which the SNARE motif interacts with the independently structured three-helix bundle, the Habc domain (13, 14), forming a four-helix bundle (Fig. 1). An additional binding site involves interactions at the outer surface of Domain 1 of Munc18-1 with a short stretch at the *N*-terminus of syntaxin-1a, the so-called *N*-peptide (14-16). The tight interaction with Munc18-1 makes syntaxin-1a unable to form a SNARE complex *in vitro* (17). An interesting feature of the closed conformation is the folding of the linker region between the Habc and H3 domains. This region folds into a small helix that lies above the four-helix bundle region that is not enclosed by Munc18-1. Two consecutive mutations, L165A and E166A, in the linker helix of syntaxin-1a allow syntaxin-1a to form a SNARE complex without breaking the interaction with Munc18-1 (14, 16, 17). This well-studied syntaxin-1a variant is called LE mutant (Syx_LE_) and is often referred to as “open syntaxin”. The LE mutant can partially rescue the defects caused by the loss of the tethering protein Unc13 in *Caenorhabditis elegans* (18, 19).

**Figure 1:**
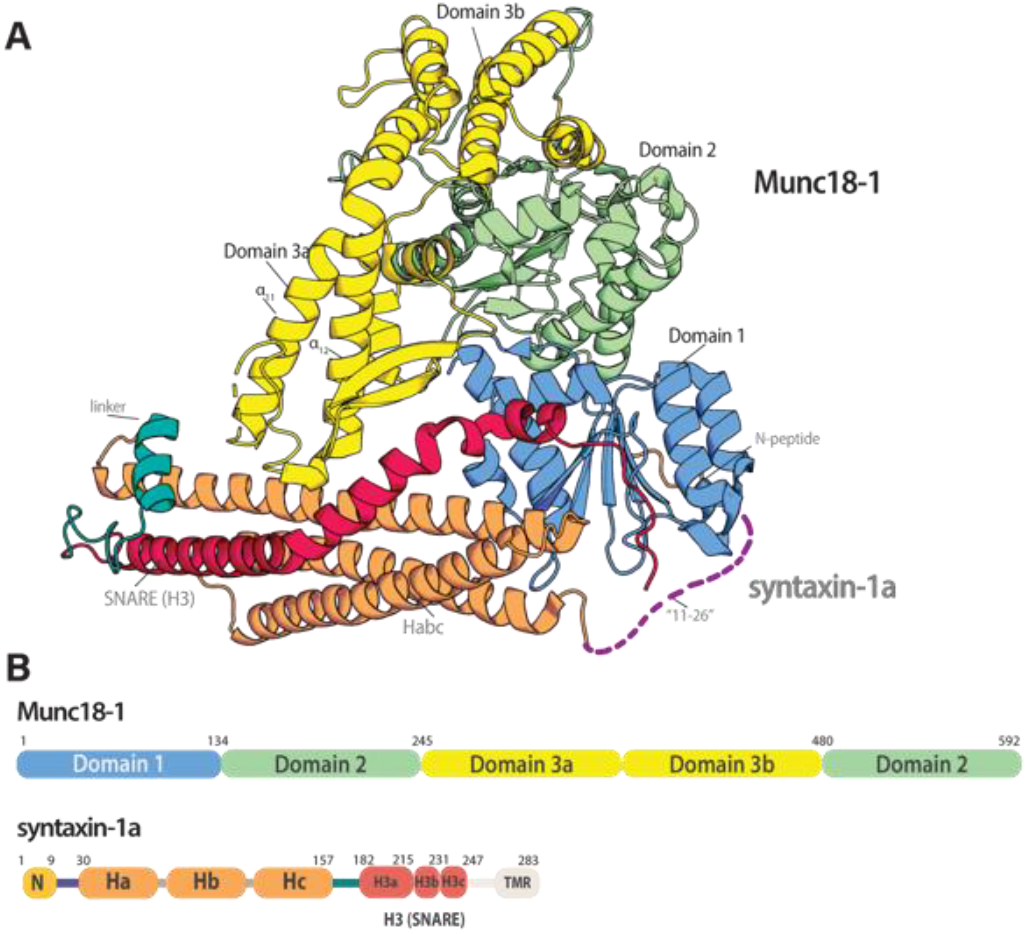
Munc18-1 forms a tight complex with syntaxin-1a. **(A)** Cartoon representation of the structure of the Munc18-1/syntaxin-1a complex ((pdb:3c98) (13, 14)). **(B)** Domain organization of the two proteins, using the same colour code as in the structure. The short *N*-peptide (N) motif, the three Habc helices, the SNARE motif (H3), and the transmembrane region (TMR) are indicated. Note that the H3 domain of syntaxin-1a, which forms an extended helix in the SNARE complex (1), has been further subdivided into Regions a, b, and c, in which bends are observed. The H3c region is buried within the central cavity of Munc18-1 (13, 14).

In individual syntaxin-1a, the linker region is flexible, allowing the protein to fluctuate between the closed conformation and a mostly unstructured open conformation (17, 20, 21). Based on this, it was originally assumed that syntaxin-1a must leave the embrace of Munc18-1 to form a SNARE complex. Syntaxin-1a would have to change its conformation so that its SNARE motif could fold into the extended helix found in the SNARE complex. *In vitro*, a similar effect was observed upon disruption of the *N*-peptide’s interactions with Munc18-1 (14). Both mutants (Syx_LE_ and Syx_ΔΝ_) still bind with high affinity to Munc18-1, but Munc18-1 appears unable to inhibit these syntaxin-1a variants from forming a SNARE complex (15). The fact that two different mutations at different sites of the complex can lead to overall similar effects suggests that syntaxin-1a remains bound to Munc18-1 and that a conformational switch within the complex might explain role of Munc18-1 in promoting SNARE complex assembly (14, 15). Indeed, it is currently thought that Munc18-1 catalyzes the formation of a SNARE complex by providing an assembly platform for the three SNARE proteins. Slowly, a clearer picture is emerging, but there are still many unanswered questions. An analysis has shown that the Munc18-1/syntaxin-1a complex can bind to SNAP-25 (22). Other studies have shown that synaptobrevin binds to the Munc18-1/syntaxin-1a complex (23-26), but how and when synaptobrevin is engaging with syntaxin-1a is unclear. It is plausible, that syntaxin-1a could already be opened up while in complex with Munc18-1 to initiate the first steps of SNARE complex formation (14).

It remains unclear how Munc18-1 facilitates the opening of bound syntaxin-1a. Individual Munc18-2 or Munc18-3 bound to the *N*-peptide of its cognate syntaxin adopt a slightly different conformation from the Munc18-1 structure from the Munc18-1/syntaxin-1a complex (27, 28). In individual Munc18-2, the helical hairpin region at the tip of domain 3a, which is formed between helices α_11_ and α_12_, adopts an extended conformation, whereas it is furled in the Munc18-1/syntaxin-1a complex. An extended conformation of the α_11_α_12_ hairpin would be sterically hindered by a syntaxin-1a in the closed conformation, suggesting that its extension could initiate the opening of the bound syntaxin-1a (29). No information structure of a Munc18-1 complex with Syx_LE_ or Syx_ΔΝ_ is available. Small structural differences were detected by Small-angle X-ray scattering between Munc18-1 in complex with Syx_wt_ and with Syx_LE_ or Syx_ΔΝ_, potentially suggesting that the variants of syntaxin-1a somewhat increased the conformational flexibility compared with the more rigid structure of the Munc18-1/syntaxin-1a_wt_ complex.

Additional insights into the putative pathway of SM protein-guided SNARE assembly have come from the study of other members of the SM protein family, which work in different trafficking steps within the cell but interact with different sets of SNARE proteins. The interaction mode of SM proteins and syntaxins (also referred to as Qa SNAREs, (30)) is evolutionarily conserved but not identical (2, 11, 31). They have probably adapted to the needs of the different transport routes, but their basic mode of operation has remained similar and may represent different stages of the reaction cascade.

The Golgi SM protein Sly1 binds tightly to the *N*-peptide region of its syntaxin partner, Sed5 (32, 33), but biochemical studies have shown that it also binds to the remainder of Sed5 and promotes its opening (34). The vacuolar SM protein Vps33 was crystalized in complex with the SNARE motif of its syntaxin partner Vam3. The SNARE motif of Vam3 is located at the same site as that of syntaxin in the Munc18-1/syntaxin-1a complex, but it is not in a closed conformation, that is, intramolecularly bound to an Habc domain. The helical hairpin region of Vps33 is extended and binds to the R-SNARE Nyv1 (35). The R-SNARE synaptobrevin has been suggested to bind to the corresponding site on Munc18-1 (24-26, 36). The structure of the SM protein Vps45 in complex with its cognate syntaxin partner, Tlg2, has been resolved recently. The helical hairpin of Vps45 is extended (29). Similar to syntaxin-1a, Tlg2 is bound by the central cavity of Vps45 but it has adopted a more open conformation, with its SNARE motif disengaged from its Habc domain, and its linker region unfolded, suggesting that Vps45 does not prevent Tlg2 from engaging with its SNARE partners. This raises the question whether the Munc18-1/syntaxin-1a complex could adopt a similar conformation when it promotes the formation of the SNARE complex. The conformational state of the Munc18-1/syntaxin-1a complex is probably controlled by other factors, for example, the CATCHR type of tethering protein Munc13, which has been shown to accelerate the transition from the Munc18-1/syntaxin-1a complex to the SNARE complex. It has therefore been speculated that a Munc13 family member can convert the Munc18-1/syntaxin-1a complex into a Vps45–Tlg2-like conformation (24, 37-39). Note, however, that the effect is only observed at micromolar concentrations of Munc13 and is much weaker than the effects of Syx_LE_ and Syx_ΔΝ_.

To gain deeper insights into the possible conformational flexibility of the Munc18-1/syntaxin-1a complex, we tested the effect of the already described and various novel point mutations and truncations on the interaction of the proteins. Our biophysical data corroborate the idea that syntaxin-1 can form a SNARE complex while bound to Munc18-1. Finally, we built a structural model of the Munc18-1/syntaxin-1a complex in a partly open conformation, based on the new Vps45/Tlg2 structure (29). The model’s structure is likely to represent an intermediate step of the reaction cascade from syntaxin-1a that is tightly bound by Munc18-1, to a more loosely bound syntaxin-1a that can subsequently begin to form a complex with its SNARE partners.

## Results

### The syntaxin-1a linker region is necessary for Munc18-1’s inhibition of SNARE assembly

In the Munc18-1/syntaxin-1a complex, the syntaxin-1a linker helix hovers over the Hc and H3 helices, sterically shielding the portion of syntaxin-1a’s SNARE motif, which is key for initiating SNARE assembly (40). We noticed that the entire linker region between the *C*-terminal end of the Hc helix and the *N*-terminal start of the H3 helix (residues 157-189) is not in direct contact with Munc18-1, and its position is mainly fixed through various contacts with residues of the Hc and H3 helices. Hydrophobic contacts, (e.g., between the conserved residues M168 and F177) within the folded syntaxin linker stabilize the position of the linker helix (Fig. 2A).

**Figure 2:**
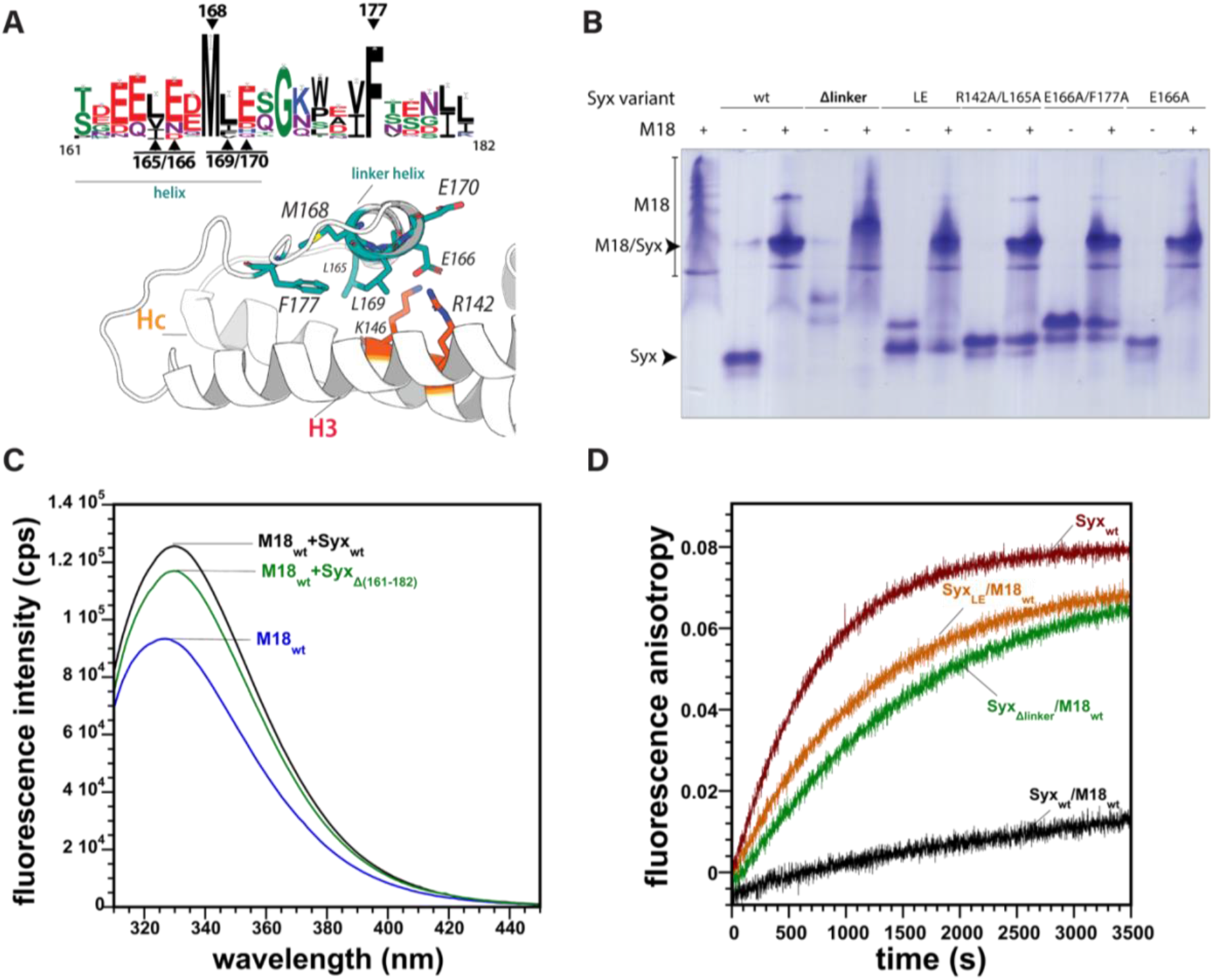
The linker region of syntaxin-1a is not essential for the interaction with Munc18-1. **(A)** The linker region of syntaxin-1a forms a short helix that interacts mostly with the H3- and Hc-helices. Key residues mutated in the study are shown as sticks. On top, the conservation of the linker region is shown as a Weblogo representation (74). Highly conserved residues are indicated. **(B)** Syntaxin-1a linker mutants form a stable complex with Munc18-1, as shown by native gel electrophoresis. Equimolar concentrations (100 μmol) of Munc18-1 and syntaxin-1a mutants were loaded individually or mixed as indicated. The band of the Munc18-1/syntaxin-1a complex is indicated by an arrow. Note that Munc18-1 by itself does not form a defined band. **(C)** Addition of syntaxin-1a to Munc18-1 leads to an increase in the tryptophan-emitted fluorescence. The emission spectra of Munc18-1 alone, or in complex with Syx_wt_ or Syx_ΔLinker_, were recorded upon excitation at 295 nm. **(D)** Ternary SNARE complex formation for several syntaxin-1a linker variants in the presence of M18_wt_, as observed by fluorescence anisotropy. Here, 40 nM of synaptobrevin labelled with Oregon Green at Cys_28_ were mixed with 500 nM syntaxin-1a, preincubated with 750 nM Munc18-1_wt_. SNARE complex formation was followed by an increase in fluorescence anisotropy upon the addition of 750 nM SNAP-25. Fluorescence anisotropy was monitored at a wavelength of 524 nm upon excitation at 496 nm. When Syx_wt_ was preincubated with Munc18-1, the SNARE complex formed slowly (black trace). Different Syntaxin-1a linker mutants (Syx_LE_: red, Syx_Δlinker_: green) were able to form a SNARE complex more rapidly than Syx_wt_ in the presence of Munc18-1. Note that we also mutated another highly conserved residue of the hydrophobic network of the linker region, F177 (Syx_F177A_). This single point mutation also somewhat released the inhibition of Munc18-1 (Fig. S1B). A double mutant, Syx_R142A_L165A_, which interfered with the polar and the hydrophobic networks on both side of the linker helix, was somewhat more disruptive than the individual exchanges (Fig. S1C).

To investigate the importance of the syntaxin-1a linker helix on the interaction with Munc18-1, we first removed the entire linker region, i.e., residues 161 to 182 (Syx_ΔLinker_). Syx_Δlinker_ formed a stable complex with Munc18-1 as assessed by native gel electrophoresis (Fig. 2B) and an increase in the intrinsic tryptophan fluorescence upon mixing of the two proteins (Fig. 2C), suggesting that the entire linker region of syntaxin-1a is not essential for tight interaction of the two proteins. This corroborates our earlier investigation that had shown that the H3 helix and the remainder of syntaxin-1a can both bind to Munc18-1 as a split syntaxin-1a (14). This has been recently confirmed by others (41, 42). When Syx_ΔLinker_ was mixed with SNAP-25 and fluorescently labelled synaptobrevin (Syb*^28^), an increase in fluorescence anisotropy was observed, showing that Syx_ΔLinker_ forms a SNARE complex. This fluorescence-based kinetic approach to following the formation of SNARE complex was instrumental to describing the inhibitory effect of Munc18-1 (14, 43). When we added Munc18-1 to the reaction, the SNARE complex formation of Syx_ΔLinker_ was slowed down somewhat, but much less than that of Syx_wt_ (Fig. 2D).

Overall, the effect of Munc18-1 on Syx_ΔLinker_ was more similar to the effect on Syx_LE_ (Fig. 2D) shown in earlier studies (14, 17). The Syx_LE_ variant carries two sequential mutations on the linker helix, L165A and E166A. How these point mutations affect the accessibility and conformation of the bound syntaxin-1a has remained unclear. Both residues form interactions within the linker. L165 is part of the hydrophobic interaction network of the linker mentioned above, forming electrostatic interactions with K146, N150, T160, and L169, as well as hydrogen bonds with T161, S162, and M168. In contrast, E166 forms ionic bridges with residues on the Hc helix (R142 & K146), while also interacting through hydrogen bonds with residues of the linker region (L169, S162, E163, E170). R142, the residue that interacts with E166, is also involved in an electrostatic interaction with R315 at the tip of Helix 11 of the helical hairpin region of Munc18-1’s Domain 3a.

The linker helix forms an obstacle that could regulate the access of the SNARE interacting partners to the *N*-terminus of syntaxin-1a’s SNARE motif. We introduced two different single helix breaking glycine mutations L165G (Syx_L165G_) and M168G (Syx_M168G_) to test whether the integrity of the linker helix plays a role in the inhibitory role of Munc18. Whereas the M168G mutation removed the inhibition only partly, the L165G mutation was almost as effective as Syx_LE_ (Fig. S1A). In the presence of Munc18-1, Syx_M168G_ assembled into a SNARE complex faster than the less disruptive variant, Syx_M168A_. This supports the idea that the integrity of the linker helix is important for the inhibition by Munc18-1.

Mutation of an adjacent pair of LE residues on the linker helix (L169A, E170A) was reported to have similar effects to Syx_LE_ (38). Indeed, the SNARE complex assembly rate of Syx_L169A_E170A_, which was very similar to that of Syx_LE_, was almost unaffected by Munc18-1 (Fig. S1B). These two residues have a similar arrangement of Syx_LE_; whereas L169 is part of the hydrophobic network within the linker, E170 is forming electrostatic interactions.

### Intrinsic fluorescence between Munc18 W28 and syntaxin-1a F34 as an indicator of a conformational change in the complex

As mentioned above, the intrinsic fluorescence of Munc18-1 increases upon binding to syntaxin-1a (14). We inspected the structure of the complex for the molecular correlate to examine this effect. We noted that the exposed tryptophan 28 (W28) on the inner surface of Domain 1 of Munc18-1 becomes buried upon binding and is then in direct contact with the phenylalanine 34 (F34) of the Ha helix of syntaxin-1a (Fig. 3A). These two highly conserved residues are plausible candidates for the fluorescence dequenching effect observed. To test this, we substituted both residues with alanines (Syx_F34A_ & M18_W28A_). Indeed, the interaction of M18_wt_ with Syx_F34A_ led to a drastic reduction in tryptophan dequenching. As expected, the intrinsic fluorescence of M18_W28A_ was reduced compared with that of M18_wt_, and almost no increase was observed when Syx_wt_ was added (Fig. S3). The proximity of W28 to F34 suggests that these residues also contribute to the stability of the complex. To test whether this interaction affects the ability of Munc18-1-bound syntaxin-1a to form a SNARE complex, we monitored the speed of SNARE complex formation after preassembly of the Munc18-1/syntaxin-1a complex. Both point mutations eased the transition from the Munc18-1/syntaxin-1a complex to the SNARE complex. The effect of Syx_F34A_ was stronger than that of Munc18-1_W28A_ (Fig. 3B). This suggests that this interaction, which occurs far from the linker region of syntaxin-1a, is also involved in maintaining the tight grip on syntaxin-1a of Munc18-1. Earlier studies showed that the fluorescence dequenching effect was somewhat diminished when Munc18-1 was mixed the Syx1a_LE_ (14). Here, we observed a somewhat lower tryptophan fluorescence increase when Munc18-1 was mixed with Syx_ΔLinker_ than that of the mix of the two wild-type proteins (Fig. 2C). Similarly, the emission maximum was lower upon mixing Munc18-1 with other linker mutants compared to that of the interaction with Syx_wt_ (Fig. S4).

**Figure 3:**
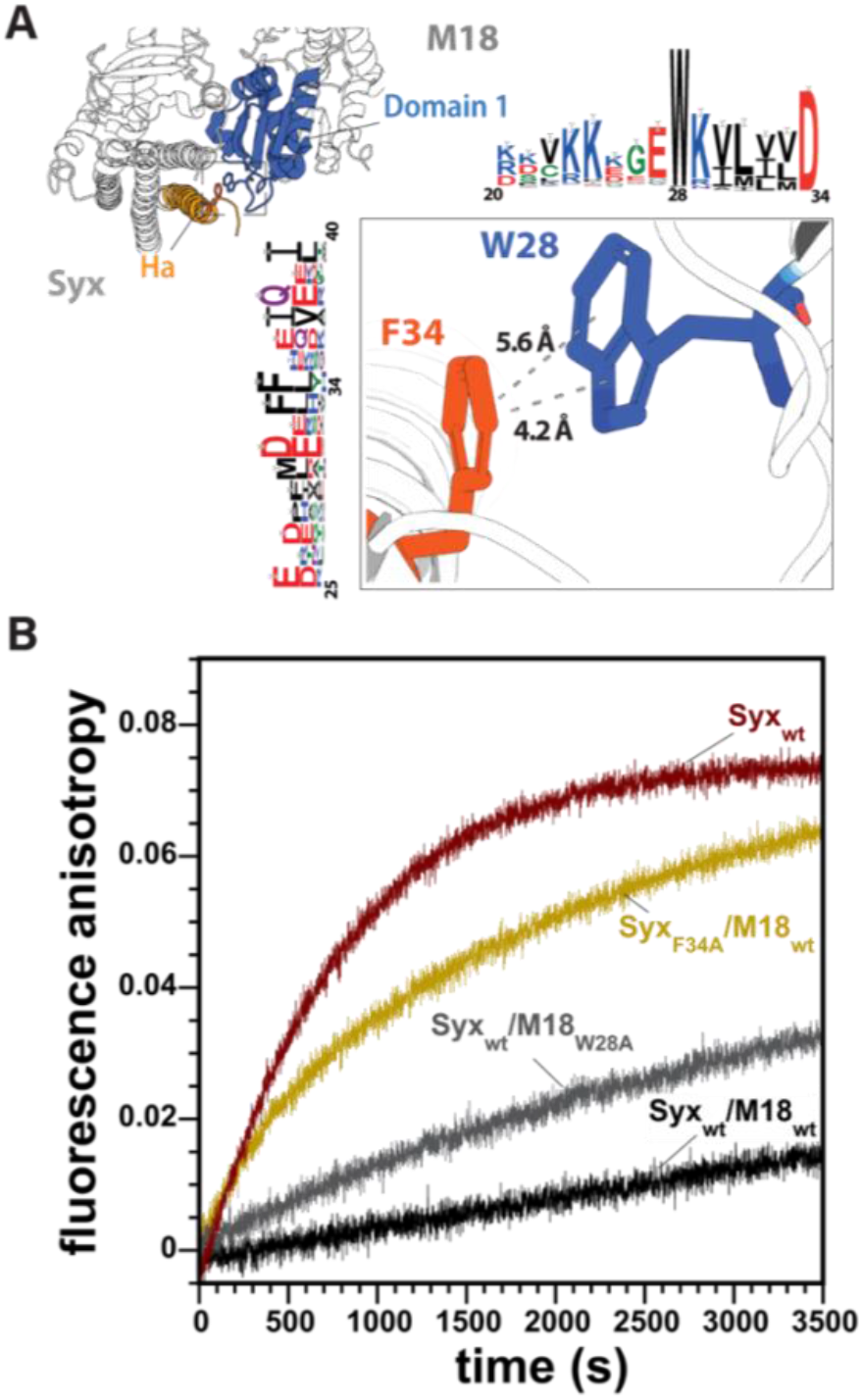
Munc18-1 W_28_ interacts through pi–pi stacking with syntaxin-1a F_34_. **(A)** In the Munc18-1/syntaxin-1a complex, the aromatic rings of syntaxin-1a F34 and Munc18-1 W28 lie within ∼5 Å d and are oriented at a dihedral angle of ∼30°. Both W28 and F34 are highly conserved across vertebrates, as shown by the Weblogo representations. (**B)** Mutations of both residues to alanines led to faster formation of the SNARE complex. Mixing experiments were carried out as described in the legend of Fig. 2.

The observed similarity in tryptophan emission changes suggests that these linker mutations lead to very similar, but still unknown, structural changes in the Munc18-1/syntaxin-1a complex. These changes, we believe,, somewhat loosens the grip on the closed conformation of syntaxin-1a by Munc18-1. In addition, the small reduction in the dequenching effect by different linker variants compared to that of syntaxin-1a_wt_ could reflect a change in the local environment of the region around W28 and F34; i.e. the conformational change in the linker region of syntaxin-1a is “sensed” by the remote interaction of W28 and F34.

### Domain 1 is clamped by the *N*-peptide and the Habc domain of syntaxin

In the structure of the complex, it can be seen that the interaction between W28 and F34 is part of an interaction network between the inner surface of Munc18-1’s Domain 1 and the Ha- and Hc-helices of syntaxin-1a (13). Note that Domain 1 contributes the majority of the interaction surface between Munc18-1 and syntaxin-1a (13). In the Munc18-1/syntaxin-1a complex, Domain 1 is in a slightly different position than in the uncomplexed Munc18-1. It probably undergoes a rotational movement during complex formation that is facilitated by a hinge region between Domains 1 and 2 (44). The binding site of the syntaxin-1a *N*-peptide is located on the outer surface of Domain 1. We wondered whether this second binding site stabilizes the position of domain 1. We noticed that the Habc domain of syntaxin-1a is connected to the *N*-peptide by a short stretch (Fig. 4A). This stretch is not visible in the structure of the complex, probably because of its high flexibility (Fig. 4A). In syntaxin-1a, it is ≈ 15 amino acids long and contains a conserved stretch of negatively charged residues. The length of the linker is conserved in other secretory syntaxins (Qa.IV (30)) as well, but their sequences vary somewhat. In order to test whether the linker clamps domain 1, we have lengthened the linker region twofold by adding the entire 15 amino acid linker (aa11-26) two more times between the *N*-peptide and the Habc domain, resulting in a syntaxin-1a variant with a linker 45aa long (Syx_3x(11-26)_). As a control, we removed the linker region, which should make it impossible for the *N*-peptide to reach its binding site.

**Figure 4:**
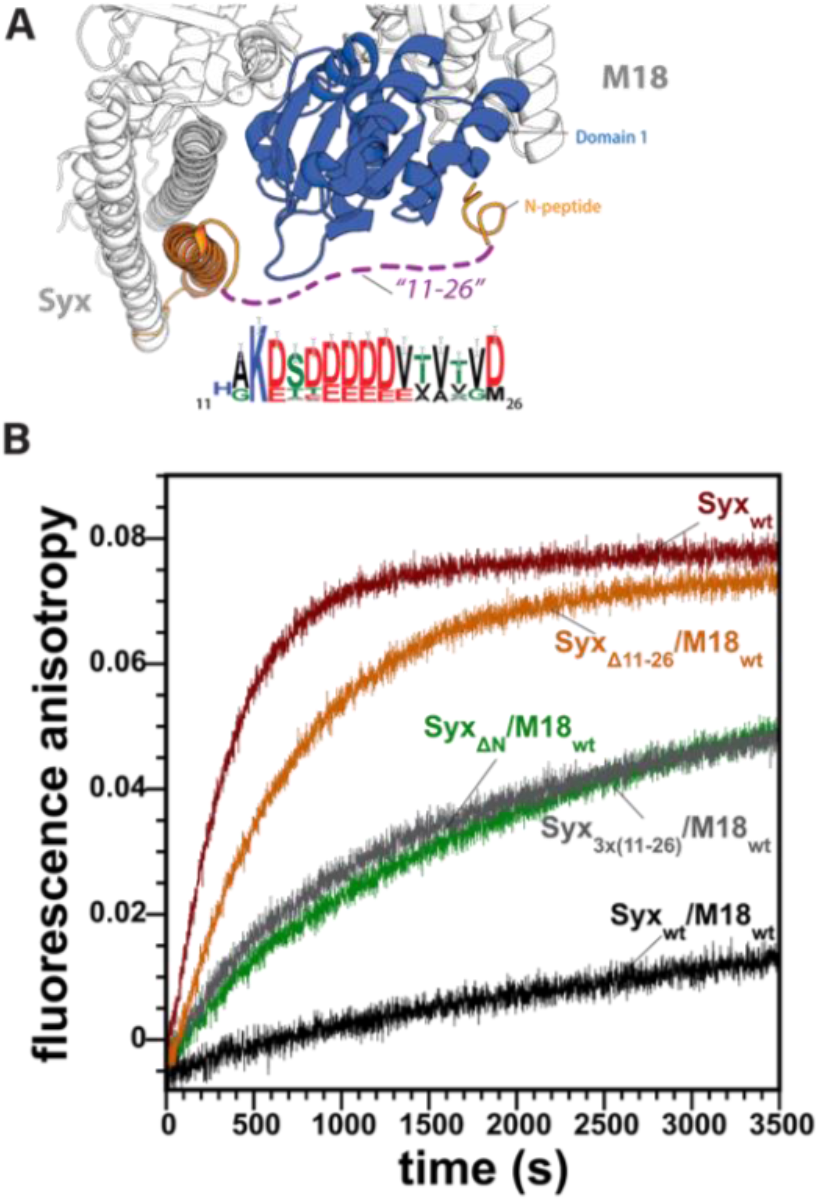
The *N*-peptide is tethered to the Habc domain by a short stretch. **(A)** Weblogo representation of the short stretch that links the Habc domain to the *N*-peptide of syntaxin 1a. Above, the section of the structure shows the position of this short region in the Munc18-1/Syntaxin-1a complex. The stretch is depicted as a dashed line, as its structure has not been resolved (13, 14). **(B)** In the presence of Munc18-1, Syx_ΔΝ_ (green), Syx_3x(11-26)_ (grey), and Syx_Δ(11-26)_ (orange) formed a SNARE complex more rapidly than Syx_wt_ (black). Mixing experiments were carried out as described in the legend of Fig. 2.

Both linker variants (Syx_Δ11-26_ & Syx_3x(11-26)_) formed a stable complex with M18_wt_ (Fig. S5). We then used our fluorescence anisotropy approach to assess the degree to which the SNARE assembly of these variants was inhibited by Munc18-1. When synaptobrevin and SNAP-25 were added to a mix of Munc18-1 and syntaxin, both syntaxin-1a variants were able to form SNARE complexes faster than Syx_wt_. The rate of Syx_3x(11-26)_ was comparable with that of syntaxin-1a without the *N*-peptide, Syx_ΔN_ (Fig. 4B). Notably, the variant with a deleted linker region, Syx_Δ11-26_, assembled into the SNARE complex even faster than Syx_3x(11-26)_ into in the presence of Munc18-1.

The transition from the Munc18-1/syntaxin-1a complex to the SNARE complex was reported to be accelerated by Munc13 (19, 37-39, 42, 45). When we added the MUN domain to our fluorescence-based SNARE complex assay, we did not observe a significant effect on the SNARE assembly reaction in the absence or presence of Munc18-1 even at high concentrations of the MUN domain. When we used SDS resistance as an indicator (8), a minor increase in SNARE complex formation in the presence of the MUN domain was detectable, but the effect was rather small in comparison with the effect of the LE mutant and the other mutants studied here (Fig. S6).

### Impact of structural changes in Domain 3a of Munc18-1 on the bound syntaxin

As mentioned above, an unfurled helical hairpin in Domain 3a of Munc18-1 is probably incompatible with binding to closed syntaxin-1a. The conformational change from a furled to an extended hairpin would have a direct impact on the four-helix bundle formed by the Habc domain and the H3a region (aa 188-211) of syntaxin-1a. In particular, the start of the H3b region (aa 212-224), which bends away from the canonical four-helix bundle, would be affected directly by an extended hairpin region. The putative conformational change in that region has been extensively studied (46-52). Several point or deletion mutations at the helical hairpin have been introduced to study the impact of this putative extension, including P335A (M18_P335A_). The highly conserved P335 is located at the tip of Helix 12 (Fig. 5A), where it forms a hinge point for the conformational transition. In order to study the effects of the confirmational transition, we utilized the previously described P335A mutation, which has been designed to artificially lock Helix 12 in an extended conformation (23, 51).

**Figure 5:**
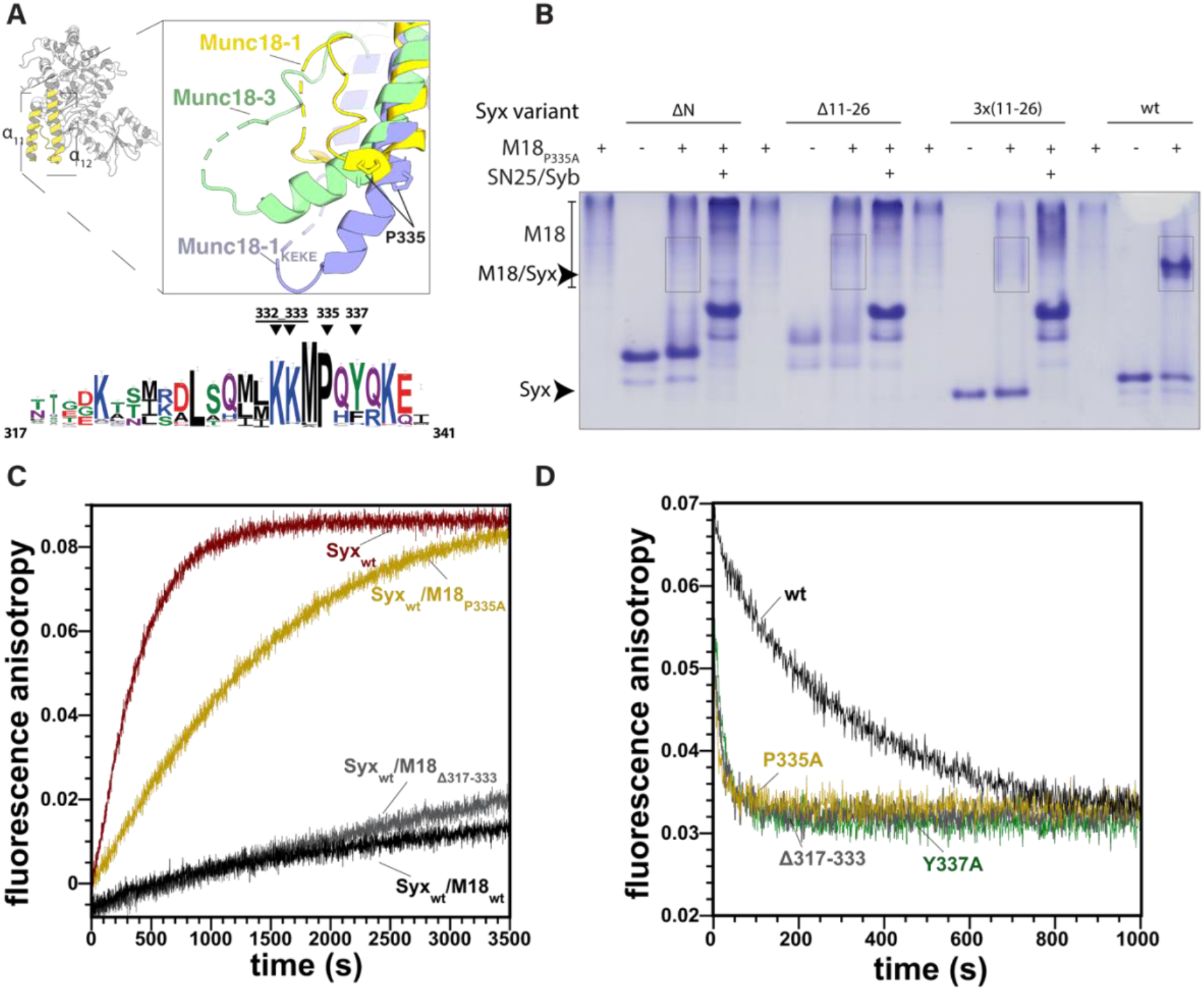
Munc18 α_11_α_12_ loop residues contribute to inhibition. **(A)** Different conformations of the α_11_α_12_ helical hairpin of Domain 3a, shown by an overlay of the Munc18-1/syntaxin-1a complex (pdb: 3c98, yellow), Munc18-3 alone (pdb: 3puk, green), and Munc18-1_K332E_K333E_ (pdb: 6lpc, violet) (top) (13, 14, 42, 48). In the Munc18-1/syntaxin-1a complex, the region of Residues 317–333 is folded upwards towards the α_11_α_12_ helices. Note, that the stretch between Residues 317–323 was not resolved. In Munc18 alone, the α_11_α_12_ loop adopts an unfurled conformation with an extension of α_12_. The very conserved proline at the tip of α12 is thought to be the hinge point for the unfurling of the helix (bottom). The conservation of the Munc18-1 helical hairpin loop is shown by a Weblogo representation. Mutated residues are indicated by arrows. **(B)** Complex formation of M18_P335A_ (55 μmol) with Syx_wt_ and different syntaxin-1a variants (34 μmol) in the absence and presence of the SNARE partners SNAP-25 and synaptobrevin (each 90 μmol), monitored by native gel electrophoresis. The positions of the monomeric proteins and complexes are indicated by arrows. Note that synaptobrevin cannot be detected in the nondenaturing gel because of its isoelectric point. Although a clear complex band was visible for the M18_P335A_/Syx_wt_ complex, the complexes of M18_P335A_/Syx_wt_ complex appeared as more diffuse bands, suggesting that these interactions were less stable. **(C)** Deletion of the hairpin loop, Munc18-1_Δ317-333,_ had only a very small effect on Munc18-1’s ability to inhibit the formation of the SNARE complex, as monitored by fluorescence anisotropy. By contrast, the mutation of the conserved P335 to an alanine rendered Munc18-1 much less able to control the complex formation of the bound syntaxin-1a, in agreement with previous studies (23, 42, 47, 51, 54). Mixing experiments were carried out as described in the legend of Fig. 2. **(D)** Determination of the off-rate of the Munc18-1/syntaxin-1a complex by competitive dissociation. An excess of unlabelled Syx1a variants (5 μM) was added to a premix of 100 nM Oregon Green-labelled Syx1a variants using the single cysteine introduced at Position 1 along with 250 nM Munc18-1; and the decrease in fluorescence anisotropy. was measured. The dissociation was fitted by a single exponential. Dissociation was faster for the Domain 3a variants (M18_P335A,_ M18_Δ(317-333),_ M18_Y337A_) than for M18wt. The dissociation rates are given in Table S1.

In agreement with earlier findings, M18_P335A_ was able to form a stable complex with Syx_wt_ that did not fall apart upon native gel electrophoresis (Fig. 5B) or size exclusion chromatography (Fig. S7A). Again, we tested the degree to which this Munc18-1 variant inhibited the formation of SNARE complex by the bound syntaxin-1a. M18_P335A_ was clearly less efficient in inhibiting syntaxin-1a’s SNARE complex formation than the wild-type Munc18-1 (M18_wt_, Fig. 5C). This corroborates earlier studies (42) and supports the notion that an extended hairpin may somehow loosened the conformational state of the bound syntaxin-1a.

Recently, the structure of another helical hairpin variant, carrying the K332E and K333E mutations (M18_KEKE_) was resolved. This mutant was first studied together with a variety of Domain 3a mutations (46, 47). M18_KEKE_ was resolved as a homodimer where Domains 2 and 3 were packed against each other, as previously also seen for squid n-Sec1 (42, 44, 46). Upon gel filtration and native gel electrophoresis, M18_KEKE_ also formed a stable complex with syntaxin-1a (Fig. 5B). Interestingly, M18_KEKE_ inhibited the formation SNARE complex as strongly as M18_wt_ (Fig. S8A).

However, why do the two variants differ in their ability to bind and hold syntaxin? We noticed that M18_P335A_ runs at a higher molecular mass than M18_KEKE_ and M18_wt_ upon gel filtration, corroborating an earlier study (42). This suggests that only M18_P335A_ dimerized at the concentration range used, whereas M18_KEKE_ remained a monomer. Possibly, M18_KEKE_ only unfurls at the high local concentrations used for crystallization. Nevertheless, the mutations support the idea that helical hairpin region of Munc18-1 can switch its conformation.

To check whether the helical hairpin is needed for the inhibition of SNARE assembly by Munc18-1, we deleted the region undergoing the conformational change, aa 317-333. As shown earlier (50), M18_Δ317-333_ was able to bind to syntaxin-1a (Fig. 5C). Compared with Munc18-1_wt_, the variant was also only slightly less able to inhibit the formation of SNARE complex, corroborating the idea that the hairpin loop is not involved in the tight embrace of syntaxin-1a.

Residues on the tip of Domain 3a also interact directly with syntaxin-1a’s residues. To elucidate the effect of the interactions on the inhibition, we used the previously described Y337A mutation (M18_Y337A_) (46, 52). Y337 is close to P335, on the tip of Helix 12, and might be involved in electrostatic interactions with the N135 of syntaxin-1’s Hc (Fig. S2A). Y337A did not affect the inhibition, whereas syntaxin-1a’s N135D substitution had some effect on the inhibition (Fig. S2C-D). We also mutated R315 at the tip of Helix 11 to an alanine (M18_R315A_) but did not observe a significant effect on the SNARE assembly rate of Syx_wt_ (Fig. S2B).

An additional region of Domain 3a, a loop between β-sheets 10 and 11 (aa 269-275), is another possible candidate for a regulatory region. Residues of this loop are involved in an extensive network of interactions with residues of the H3 helix of syntaxin-1a (Fig. S9A). This region was previously found to be disordered in the absence of the syntaxin-1a *N*-peptide binding to Munc18-1 (15). When we tested a variant in which the loop was deleted (M18_Δ269-275_), in our fluorescence-based assay, we observed a small acceleration in SNARE complex assembly compared with M18_wt_ (Fig. S9B). However, the substitution of the E224 residue of syntaxin-1a with alanine (Syx_E224A_), which interacts with S269 of Munc18-1, had only a small effect on the inhibition (Fig. S9C). To further evaluate the effect of Munc18-1 mutations on the interaction with Syx_wt_, we used another fluorescence anisotropy-based assay. The dissociation rate of M18_variants_/Syx complexes was determined by fluorescence anisotropy decay (Fig. 5D). As expected, all mutations increased the dissociation rate to some degree (Table S1).

### Stability of the Munc18-1/syntaxin-1a complex assessed by molecular dynamics

To understand the different effects of the mutations tested, we carried out molecular dynamics (MD) simulations of the Munc18-1/syntaxin-1a complex (13, 14). We used the conformations obtained during non-restraint simulations to calculate the energy of their interaction, and the contributions to this energy of every residue in the complex by using the molecular mechanics generalized Born surface area (MM-GBSA) method. We also estimated the importance of every residue side chain for protein stability by using FoldX 5.0 alanine scan calculations.

As mentioned above, two regions of Munc18-1 contributed the most to the interaction, namely the inner cavity formed by Domains 1 and 3 (13) and the outer surface of Domain 1 (14). The most important residues for the interaction with syntaxin1a found on Munc18-1 are the hydrophobic P335 and I271, and the charged R64 on the *N*-terminus, whereas those contributing the most to the energy of interaction on syntaxin1a are residues of the H3 domain: I233 and N236. The contribution of the most important residues in these regions are highlighted in Fig. S11 and listed in Table S2.

We also took a closer look at the areas where we had introduced mutations. Very few residues in the helical linker region between the Habc domain and the H3 domain of syntaxin, namely E166, D167, and E170, contribute moderately (below 2 kcal/mol) to the interaction with Munc18-1. The R315 of Munc18-1 seems to be their only interacting partner, contributing 3.4 kcal/mol to the interaction’s energy. The highly conserved F177 contributes 3.6 kcal/mol to the conformational stability of the linker region, as estimated by FoldX. Similarly, the hydrophobic side chains of L169 and L165 of syntaxin-1a contribute to stability of the linker region (2.0 kcal/mol and 1.6 kcal/mol, respectively), whereas the conserved M168 contributes less, somewhat unexpectedly (0.5 kcal/mol).

### A model of the Munc18-1/syntaxin-1a complex in a more open conformation

Our biochemical investigations support the idea that Munc18-1-bound syntaxin-1a is capable of forming a SNARE complex. The interaction with Munc18-1 could serve to prepare syntaxin-1a for the formation of SNARE complex, which is facilitated by a slight change in the conformation of the Munc18-1/syntaxin-1a complex. We noted that the structure of a homologous complex (Vps45/Tlg2) was recently resolved in a more open conformation, in which Domain 3a’s helical hairpin loop of the SM homolog, Vps45, is was unfurled, and the SNARE domain of the syntaxin homolog, Tlg2, was extended, and appeared to be ready to engage with its SNARE partners (29). Could that structure resemble the intermediate conformational state of the syntaxin1-a/Munc18-1 complex that has remained elusive thus far?

Terefore, we generated a model of the Munc18-1/syntaxin-1a complex based on the structure of Vps45/Tlg2 structure by using Modeller 9.18 software (53) (Fig. 6). In this model, the hairpin loop of Domain 3a of Munc18-1 is in an unfurled conformation. The β_10_β_11_ loop is changed as well: the β_10_ and β_11_ sheets are shorter and have moved away somewhat from the H3b region of syntaxin. The *N*-terminal region of the H3 domain, which stretches slightly beyond the Hc and H3b region, is now in an extended α-helical conformation that does not form a four-helix bundle with the Habc domain but follows the extended Helix 12 for a short stretch. The SNARE layer residues of the free α-helix, formed by the H3a and H3b region, are twisted towards the outer surface and appear to be ready to bind to other SNAREs. The H3c region is almost unchanged in the binding cavity between the Domain 1 and Domain 3 of Munc18-1. Similarly, the *N*-peptide region remains bound to the outer surface of Domain 1 in the model’s structure. Both interaction surfaces contribute substantially to the interaction energy, supporting the notion that they can remain bound in the Munc18-1/syntaxin-1a complex in the more open conformation.

**Figure 6:**
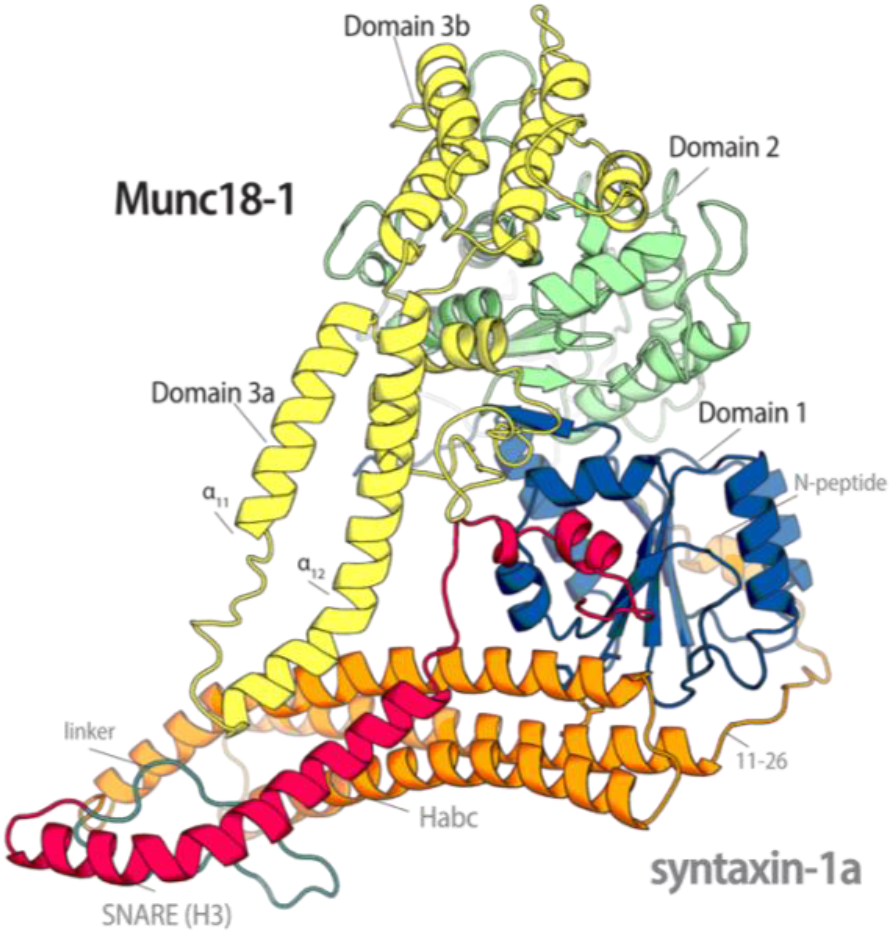
Model of Munc18-1/Syntaxin-1a in an intermediate conformation. Cartoon representation of the Munc18-1/syntaxin-1a complex modeled by using the Vps45/Tlg2 structure as a template.

The three-helix bundle formed by the Habc domain remained intact, but one side of the Habc domain rotated away by ∼30 degrees from its position in the crystal structure, following the unfurled hairpin helix of Domain 3a of Munc18-1. The region of the Habc domain that points into the interior of the binding cavity barely changed its position. This can be seen by the small change in the interaction between W28 and F34 (Fig. S10), for example. The entire linker region between the Habc and H3 domain was modeled in an unstructured conformation and moved away. The model is consistent with our biochemical findings, which suggested a small conformational change in the Munc18-1/syntaxin-1a complex. The model highlights the structural changes that take place to prepare syntaxin-1a for the upcoming formation of the SNARE complex, while syntaxin-1a remains in association with Munc18-1. The conformational changes occurred at multiple sites in the complex. It is unclear, however, how they are triggered and in what order they occur.

## Discussion

The SM protein Munc18-1 is an essential component of the apparatus of neurotransmitter release. It is thought to provide a folding platform for the formation of SNARE complex. A picture emerges in which Munc18-1 first binds syntaxin-1a in a closed conformation, preventing the bound syntaxin-1a from interacting with its SNARE partners. This allows precise control of the important neurotransmitter release reaction. Many studies in recent years have suggested that a structural change in the Munc18-1/syntaxin-1a complex subsequently occurs, which drives the reaction further. The details of this change have been slow to emerge, and we have attempted to track down this elusive step in the present study.

Originally, it was thought that the LE mutations in the linker region of syntaxin-1a prevented binding to Munc18-1. However, it was then shown that this mutation only slightly reduced affinity, but Munc18-1 lost control of the bound syntaxin-1a through these point mutations; the LE mutant could form a SNARE complex despite tight binding to Munc18-1. The slight difference in the change in intrinsic tryptophan fluorescence when it formed a complex with the LE mutant rather than wild-type syntaxin-1a indicated a subtle conformational change in the complex (14). This was corroborated by small-angle X-ray scattering measurements (15). However, no high-resolution structure of the complex of Munc18 with the LE mutant is available.

We have shown here that the entire linker region of syntaxin-1a plays only a small role in its interaction with Munc18-1. In fact, it can be excised altogether without greatly altering their binary interaction. Our data corroborate the notion that the linker region acts as a kind of shield over the SNARE domain of syntaxin-1a. The folding of the linker is stabilized by intramolecular, hydrophobic, and hydrophilic interactions. Interference with these networks led to a total, or partial, loosening of Munc18-1’s hold, depending on how greatly the linker region was destabilized by the different mutations.

Several other mutations, including the removal of the *N*-peptide of syntaxin-1a, led to very similar results, i.e. a reduced change tryptophan fluorescence and the loss of inhibition, suggesting that comparable conformational changes were induced, although the affected regions were on different ends of the complex. Is there a structural connection, as suggested earlier (15)? We identified the inbuilt fluorescence sensor as the highly conserved W28 on the inner surface of Munc18-1. In the complex, W28 is in close proximity to F34 on the surface of the Ha helix of syntaxin-1a. We found that W28 is also an important contributor to the stability of the interaction. The *N*-peptide binding site is on outer side of Domain 1, whereas W28 interacts with its inner surface. Accordingly, it seems that Domain 1 is held in place by the two interaction sites. We tested this idea by extending the stretch that connects the *N*-peptide to the Habc domain, and again observed that the grip of Munc18-1 was loosened. It could be that the whole syntaxin-1a bundle, the Habc domain, and the H3a region, can move somewhat in the complex when weakening mutations are introduced to different interaction surfaces. However, these mutations do not greatly affect the tight clasp of the H3c stretch within the cavity of Munc18-1.

A slight movement of syntaxin-1a could lead to changes in the structure of Munc18-1 or vice versa. A small rotational movement of Domain 1 was found to be different between isolated Munc18 and Munc18 in a complex with syntaxin (11). The other structural difference between these structures is the extension of the hairpin loop of Domain 3a in uncomplexed Munc18s. The extension of Helix 12 would sterically hinder the binding of syntaxin in a tightly closed conformation. The P335A mutation most probably locks helix 12 in the extended position. Nevertheless, a stable complex with syntaxin can form. Again, syntaxin does not appear to be bound in the tightly closed conformation, as Munc18 can barely prevent the formation of the SNARE complex. It should be noted that the corresponding P335A mutation in *C. elegans* (P334A) partially rescued the impairments in neurotransmitter release caused by the deletion of its Munc13 homolog, Unc13 (54). This is very similar to the effect of the LE mutation (19), suggesting that both mutations induce a similar change in the Munc18/syntaxin complex. What conformation does the Munc18/syntaxin complex adopt? How can we visualized how syntaxin remains stably bound to Munc18-1 even though the interaction with its SNARE partners is now possible? Since it has not yet been possible to obtain a high-resolution structure of this elusive state yet, we used the recently published structure of the distantly related Vps45/Tlg2 complex to create a model of the Munc18-1/syntaxin-1a complex in a more open conformation. The modelled structure (Fig. 6) is very similar to that Vps45/Tlg2 complex. Compared with the X-ray structure of the Munc18-1/syntaxin-1a complex (13, 14), the SNARE-motif of syntaxin-1a is no longer in association with its Habc domain in the modelled structure. Moreover, the linker region is unfolded, and the entire N-terminal region of the SNARE-motif forms a helix that appears to be ready to interact with the other SNARE-helices. It needs to be kept in mind that our model needs further experimental validation. In the complex, resolved by X-ray or in our model, Munc18-1 interacts with almost the entire syntaxin-1a molecule. Syntaxin-1a, appears to be very flexible and can adopt different structures (1, 13, 14, 55, 56). *In vivo* studies have confirmed that all regions of syntaxin play a role in the release of neurotransmitter (16). Our model provides a visualization of the probable intermediate conformation during the transition from a closed conformation to a looser conformation of syntaxin-1a while in a complex with Munc18-1. The exact sequence of the conformational change that affects different regions of the complex still needs to be clarified.

Very probably, to switch from the closed conformation to the more open conformation of the Munc18-1/syntaxin-1a complex, an energy barrier must be overcome. This energy barrier permits tight regulation by other factors. The best candidate for this regulatory role is the protein Munc13. Several publications have shown that the addition of its central MUN domain accelerates SNARE protein-driven membrane fusion (38, 57-59). We tested the MUN domain in our assays but detected only minor accelerating activity. Acceleration required a high concentration of the protein, and the measured acceleration was lower than that of the LE mutant, corroborating the observations by others. It should be noted, however, that our approach only measured the speed of SNARE complex formation in solution. It could be that Munc13 activity is only measurable in the presence of membranes, i.e., when the SNARE proteins are anchored in the membrane. It has been postulated that Munc13 interacts with complex close to the syntaxin-1a linker/Munc18-1 Domain 3a region, which represents a potential target site where syntaxin-1a could be opened (24, 37-39, 42, 59). Our study demonstrated that slight destabilization of the syntaxin-1a linker’s interactions affected the closed conformation of syntaxin-1a. It is possible that other interactions of the linker residues with Munc13 residues could be the trigger to initiate the release of opening. For example, the interactions of the highly conserved F177 and M168 in the syntaxin-1a linker region could potentially be a target for the conserved aromate-rich NF region on the surface of Helix H6 in Subdomain B of the MUN domain (38, 60).

Other points of attack for Munc13 or other controlling factors would be Domains 1 and 3a. So far, Domain 1 has been rather neglected, but our experiments suggest that the position of domain 1 plays a role in keeping syntaxin-1a bound in the closed conformation; the *N*-peptide of syntaxin-1a binds on the opposite side of Domain 1 as if it were tightening its position. Domain 3a has a built-in mechanism, a kind of lever arm, that can elongate, which is incompatible with binding to the closed conformation of syntaxin-1a. However, it is unclear whether the extension of Helix 12 is active or if it responds to an earlier opening of syntaxin. In our model, the extended Helix 12 acts as a guardrail that directs the H3 domain into a specific position. The extended helix 12 also allows synaptobrevin to bind to the hairpin region, as shown by a recent study (36). It appears as if synaptobrevin binds to Munc18-1 in such a way that is positioned for a subsequent interaction with the H3 domain. It should be noted that synaptobrevin binds with low affinity to Munc18-1, and to study its structure, synaptobrevin had to be cross-linked to syntaxin (61). The idea that the R-SNARE binds to Munc18 was first inspired by the crystal structure of the SM protein Vps33, which binds both the H3 domain of the syntaxin Vam3 and, in another complex, the R-SNARE Nyv1 (35). It is not clear, however, whether the positioning of the R-SNARE helix represents an early or late step of the reaction cascade. Note that earlier biochemical findings suggested that the H3 domain of syntaxin interacts first with SNAP-25, possibly with the *N*-terminal SNARE motif of SNAP-25 (55, 62, 63). Do the *N-*terminal regions of the H3 domain of Munc18-bound syntaxin and SNAP-25 have to assemble first to prepare a binding site for synaptobrevin? Indeed, the question of the timing of the interaction of SNAP-25 has remained unanswered. Note that the complex of Munc18-1 with syntaxin-1a and SNAP-25 was isolated (22) a while ago but its structure is still elusive.

All the structures of the different SM protein types are almost congruent, but the interactions with the respective syntaxin partners are different. We assumed that they represent different steps of a conserved reaction cascade, but we must not forget that the proteins have gone separate ways since the time of last eukaryotic common ancestor, about 2 billion years ago. It could be that certain peculiarities can be found only for one or the other pair. Interestingly, the *C-*terminal region of the H3 domain is tightly bound in the central cavity in all three pairs, Munc18/syntaxin, Vps45/Tlg2, and Vps33/Vam3, and probably presents a common mode of interaction of SM proteins and syntaxins. Interestingly, in our model of the more open conformation of the Munc18-1/syntaxin-1a complex, the hydrophobic interactions in the cavity of the Munc18-1 Domain 3a continued to hold the rest of syntaxin, preparing it for the next reaction step. These interactions appear to be so strong that that they are probably not released spontaneously, suggesting that the next step in the cascade towards complete SNARE assembly needs additional factors. In fact, preliminary MD simulations suggested that the more open conformation of the Munc18-1/syntaxin-1a complex would need to be stabilized by other protein(s).

The release of neurotransmitters is controlled by complex machinery consisting of various proteins. The SM protein Munc18-1 occupies an important role, as it crucially regulates the interaction of the SNARE protein syntaxin-1a, most probably by providing a template for the formation of the central SNARE complex. Our biochemical studies removed this complex from the larger protein network to better characterize its reactions and conformational changes. Our data support the idea that Munc18-1 prepares syntaxin-1a for forming the SNARE complex. Ultimately, the goal is to understand the entire mechanism and its reaction cascade. New insights into the reaction events and additional intermediate conformations are already coming from the cryo-electron microscopy field (36, 64), so the exact sequence of events leading to the release of neurotransmitters will, hopefully, soon be understood in its entirety.

## Experimental Procedures

### Constructs and protein purification

All bacterial expression constructs were derived from rat (*Rattus norvegicus*) and were cloned into a pET28a vector that contains an *N*-terminal thrombin-cleavable 6xHis tag. The constructs for the SNARE proteins cysteine-free SNAP25(1–206), the soluble portion of syntaxin-1a (Syx(1–262)), syntaxin-1a lacking the *N*-peptide (Syx_ΔΝ_(25–262)), and Syx_LE_ (Syx_L165A/E166A_(1–262)), and the soluble portion of synaptobrevin-2 (Syb2(1–96)), as well as full-length Munc18-1 (Munc18-1(1–586)) have been described before (14, 20, 65). Likewise, the single-cysteine synaptobrevin (Syb_C28_) and syntaxin (Syx_M001C_) have been described (66). Codon-optimized versions of the following protein sequences were synthesized and subcloned into the pET28a vector (GenScript): Different syntaxin-1a (1– 262) variants with single– and double–point mutations were prepared: Syx_RI_(R151A/I155A), Syx_L169A/E170A_(L169A/E170A), Syx_F34A_(F34A), Syx_N135A_(N135A) Syx_N135D_(N135D), Syx_R142A_(R142A), Syx_K146A_(K146A), Syx_L165A_(L165A), Syx_L165G_(L165G), Syx_E166A_(E166A), Syx_M168A_(M168A), Syx_M168G_(M168G), Syx_E170A_(E170A), Syx_F177A_(F177A), Syx_E224A_(E224A), Syx_R142A/L165A_(R142A/L165A), and Syx_E166A/F177A_(E166A/F177A). A syntaxin-1a construct with the stretch between the *N*-peptide and Habc domain was prepared, Syx_Δ11-26_. Another deletion construct was cloned, in which the linker between the Habc domain and the SNARE motif was removed: Syx_Δlinker_(Δ161–182). In addition, a construct with an extended stretch between the *N*-peptide and Habc domain was prepared. It contained the region between aa 11–26 three times in a row, Syx_3x(11-26)_. Different variants of full-length Munc18-1(1–586) with mutations were prepared: M18_KEKE_ (K332E/K333E), M18_P335A_ (P335A), M18_Y337A_ (Y337A), M18_W28A_ (W28A), and M18_R315A_ (R315A). The following Munc18-1 deletion variants were cloned: M18_Δ317-333_ (Δ317–333)) and M18_Δ269-275_ (Δ269–275). A construct of the MUN domain was prepared as described earlier (38): MUN_933_(Munc13-1 (933–1407, EF, 1453–1531)).

All constructs were expressed in *Escherichia coli* BL21 DE3 cells. Proteins were purified by Ni^2+^ affinity chromatography, followed by ion-exchange chromatography essentially as previously described (14, 20). To avoid non-specific proteolysis of the syntaxin-1a *N*-peptide, His-tag cleavage by thrombin (14) was omitted from all protein preparations. Note that the preparations of Munc18-1 were always used fresh after purification and were not stored at -80°C, as freezing and thawing affects the function of the protein.

### Size exclusion chromatography

Size–exclusion chromatography was used to identify protein–protein interactions. Prior to separation on a Superdex 200 column, the proteins were incubated in equimolar amounts for 30 min at room temperature. Elution was achieved by washing the column with a Tris buffer (150 mM NaCl, 20 mM Tris pH 7.4, 1 mM DTT) at a flowrate of 0.3 ml/min.

### Electrophoresis

Routine SDS-PAGE was carried out as described by Laemmli (67). Nondenaturing gels were prepared and run in a manner identical to that of SDS-polyacrylamide gels except that SDS was omitted from all buffers. All gels were stained by Coomassie–Blue.

### Fluorescence spectroscopy

All measurements were carried out in a PTI QuantaMaster spectrometer in T-configuration equipped for polarization. All experiments were performed in 1-cm quartz cuvettes (Hellma) in a PBS buffer (20 mM sodium phosphate, pH 7.4; 150 mM NaCl; 1 mM DTT). Intrinsic fluorescence measurements were performed at an excitation wavelength of 295 nm. Emission spectra were recorded in the range of 310–450 nm. All spectra were corrected for background fluorescence from the buffer. Measurements of fluorescence anisotropy, which indicates the local flexibility of the labelled residue and which increases upon complex formation, were carried essentially as described by (40, 63). The single-cysteine variants, (Syx^*1^) and (Syb^*28^) were labelled with Oregon Green 488 Maleimide according to the manufacturer’s instructions (Invitrogen). Fluorescence anisotropy was recorded at λ_emi_=524 nm upon excitation at λ_exc_=496nm. The G factor was calculated according to G = I_HV_/I_HH_, where I is the fluorescence intensity, and the first subscript letter indicates the direction of the exciting light and the second subscript letter shows the direction of emitted light. The intensities of the vertically (V) and horizontally (H) polarized light emitted after excitation by vertically polarized light were measured. The anisotropy (r) was determined according to r=(I_VV_–G I_VH_)/(I_VV_ +2 G I_VH_).

### Molecular dynamics simulations

To calculate the trajectories and analyze the data from molecular dynamics (MD) simulations, we used the all-atom CHARMM36m force field parameters (68) and Gromacs 2021.3 (69) The 3c98 structure of the Munc18-1/syntaxin-1a complex (13, 14) was used as a starting point. The few missing loops were obtained from the AlphaFold2 (70) models of individual syntaxin-1a and Munc18-1. The complex was placed in a dodecahedral box filled with TIP3P water. As counterions, Na^+^ and Cl^-^ ions were added to the simulation box to neutralize the system and to reach a concentration of 0.15 M. The system was subjected to 1000, 1500, and 2000 steepest descent minimization steps, and, final conformations of each minimization were used as the starting points for the three trajectories. Each simulated system temperature was gradually increased to 300 K. The simulated annealing method was used, with three rounds consisting of linear heating by 100 K for 50 ps and subsequent MD simulation for 50 ps at the given temperature until a temperature 300 K was reached. The solute and the solvent were coupled to separated Berendsen heat baths during the procedure. The systems were subjected to an equilibration MD simulation lasting 100 ps with position restraints on the protein atoms with a force constant equal 300 kj/mol/nm^2^ at a constant volume and a temperature of 300 K. We continued equilibrating the systems for 100 ps MD simulation without position restraints at a constant volume and constant temperature of 300 K to ensure uniform water distribution. This was followed by an additional MD equilibration run of 100 ps at a constant temperature of 300 K and under constant pressure conditions of 1 bar. MD simulations of production were calculated via the NPT scheme at a constant temperature of 300 K and a constant pression of 1 bar. A Nosé–Hoover thermostat and a Parrinello–Rahman barostat were used. The electrostatic interactions were calculated by the particle mesh Ewald method. The cutoff value was 1.2 nm. The Van der Waals interactions were switched between 1.2 and 1.0 nm. The time step during the simulations was set to 2 fs. An analysis was performed on 100 frames extracted regularly from each production trajectory saved between 1 ns and 120 ns of the simulation. The contribution of the residues to the free energy of binding was estimated by the MM-GBSA method [8]. In this method, the free energy of binding in the complex is estimated using the equation: ΔG_bind_ = ⟨ΔG^0^_bind_⟩ + ⟨ΔG_desolv_⟩ - ⟨TΔS⟩, where ΔG_bind_ is the free energy of binding, ΔG^0^_bind_ is the contribution of the gas phase, ΔG_desolv_ is the desolvation energy upon binding, and TΔS is the entropic contribution. The brackets indicate that the values were averaged over the conformations derived from the MD trajectory. In the MM-GBSA approach, it is possible to decompose the total free energy of binding into the contributions of atoms or residues. In this study, we used the approach decomposition of the free energy of binding as described in (71). As we were interested in the order of residues’ contributions to the free energy of binding and not in the absolute values, we omitted the costly entropy term, being aware that the estimated absolute values of the contributions to free energy are not precise.

### Contributions to stability calculated by FoldX

The contribution of the residues to the protein’s stability was assessed by FoldX 5.0 software (72) using the crystal structures of Munc18-1 and syntaxin-1a taken from the 3c98 structure of the Munc18-1/syntaxin-1a complex (14). The stability was estimated by FoldX as the difference in free energy between the folded and unfolded protein. The contribution of each residue’s side chain was estimated as the difference in stability between the wild-type protein and the stability of protein variant with residue mutated to alanine.

## Homology modeling

A homology model of the open conformation was constructed with Modeller 9.18 (53) software. The Vps45 and Tlg2 protein complex stored under the 6xm1 code in the PDB (29) was used as a template to model the structure of the rat Munc18-1/syntaxin-1a complex. In total, 100 models were produced and the one with best DOPE score (73) was chosen for further optimization. The loops that were non-crystalized in the 6xm1 structure, were additionally modelled with Modeller’s *ab initio* loop modeling procedure. The best loop structure was chosen on the basis of the lowest DOPE score value. The Ramachandran plot of the main chain conformation of the resulting structure did not show any residues in the disallowed regions (61). The model structure can be found aat the Zenodo repository (https://doi.org/10.5281/zenodo.7354961).

## List of abbreviations

(SNAREs): <u>N</u>-ethylmaleimide-<u>s</u>ensitive factor <u>a</u>ttachment <u>re</u>ceptors
(CATCHR): <u>c</u>omplex <u>a</u>ssociated with <u>t</u>ethering <u>c</u>ontaining <u>h</u>elical <u>r</u>ods
(SAXS): <u>S</u>mall-<u>a</u>ngle <u>X</u>-ray <u>s</u>cattering
(SM): <u>S</u>ec1/<u>M</u>unc18
(MD): molecular dynamics

## Acknowledgements

This work was supported by the Swiss National Science Foundation (Grant 31003A_182732 to D.F.). We thank the Division de Calcul et Soutien à la Recherche of the UNIL for access to the university’s computer infrastructure. The molecular modeling support was provided by the Protein Modeling Unit of the University of Lausanne. We thank all members of the Fasshauer Laboratory for helpful discussions.

## Author contributions

I.S., J.I., and D.F. designed the study, performed the experiments, and analysed the data;

I.S. and D.F. wrote the paper; and all authors reviewed and commented on the manuscript.

