## supplements for "Exploring the conformational changes of the Munc18-1/syntaxin 1a complex"

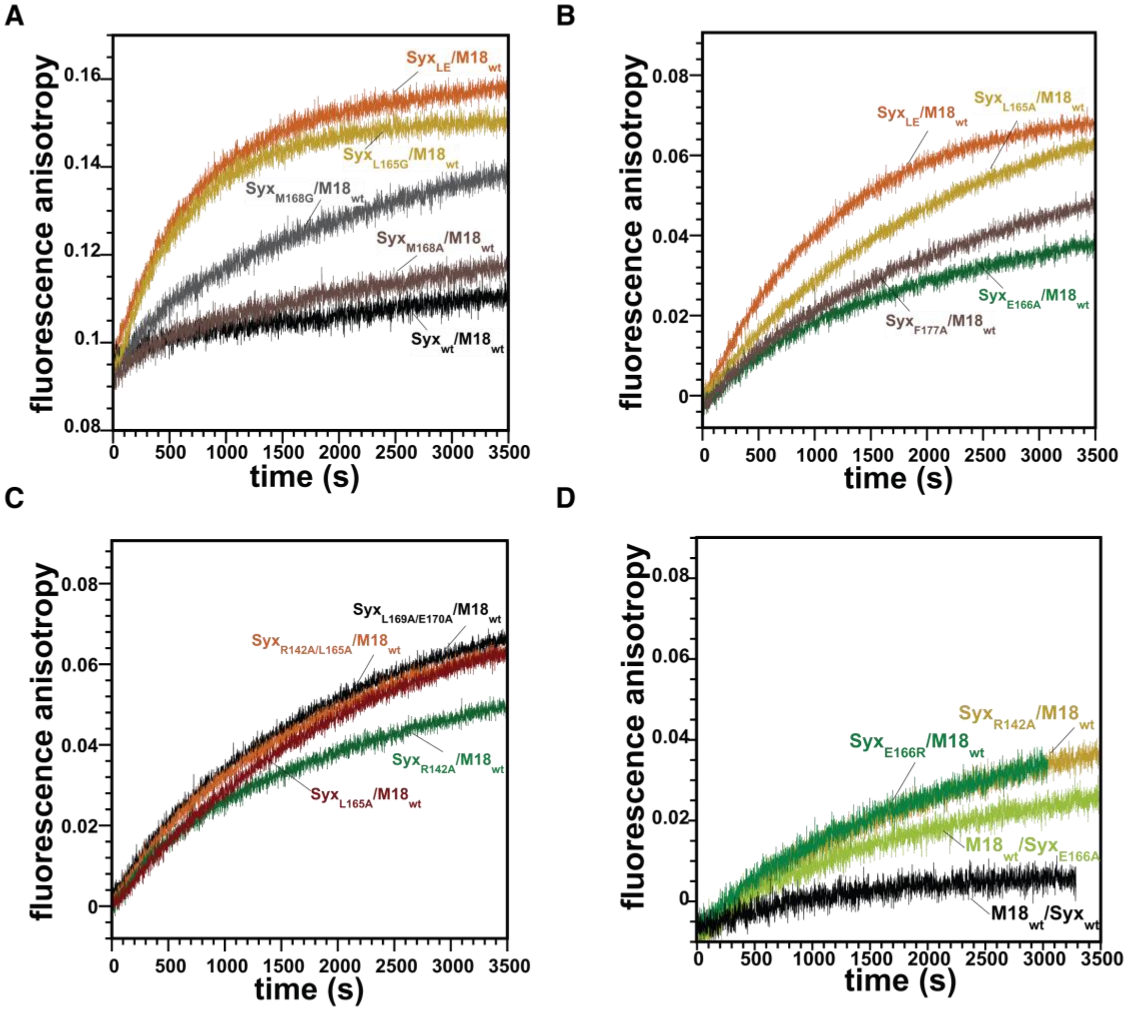


**Figure S1: The strength of the inhibitory effect of Munc18-1$_{wt}$ on the formation of the SNARE formation varies among syntaxin-1a linker mutants.**

**(A)** Syx$_{L165G}$ (ochre), carrying a helix-breaking glycine residue, forms a SNARE complex more rapidly in the presence of Munc18-1 compared with Syx$_{wt}$ (black). The effect of the single exchange is similar to that of Syx$_{LE}$ (orange; see also Fig. 2). A similar exchange at Position 168, Syx$_{M168G}$ (grey), had a less drastic effect. Its substitution to an alanine residue (Syx$_{M168A}$ ; brown) had only a small effect. **(B)** Single exchanges of the LE positions on the linker helix, Syx$_{E166A}$ (green) and Syx$_{L165}$ (ochre), were less disruptive than the double mutation in Syx$_{LE}$. Substitution of the conserved F177 to alanine (brown) also caused moderate disruption. **(C)** The syntaxin-1a double mutants Syx$_{L169A_E170A}$ (black) and Syx$_{R142A_L165A}$ (orange) assembled into SNARE complexs faster than mutants carrying a single mutation in the linker region, Syx$_{R142A}$ (green). **(D)** Mutations of E166 to an arginine, Syx$_{E166R}$ (green), and alanine (Syx$_{E166A}$) significantly reduced the inhibition of SNARE complex assembly by Munc18-1. Inhibition was also reduced when R142 was substituted by an alanine (Syx$_{R142A}$; ochre). All mixing experiments were carried out as described in the legend of Fig. 2.

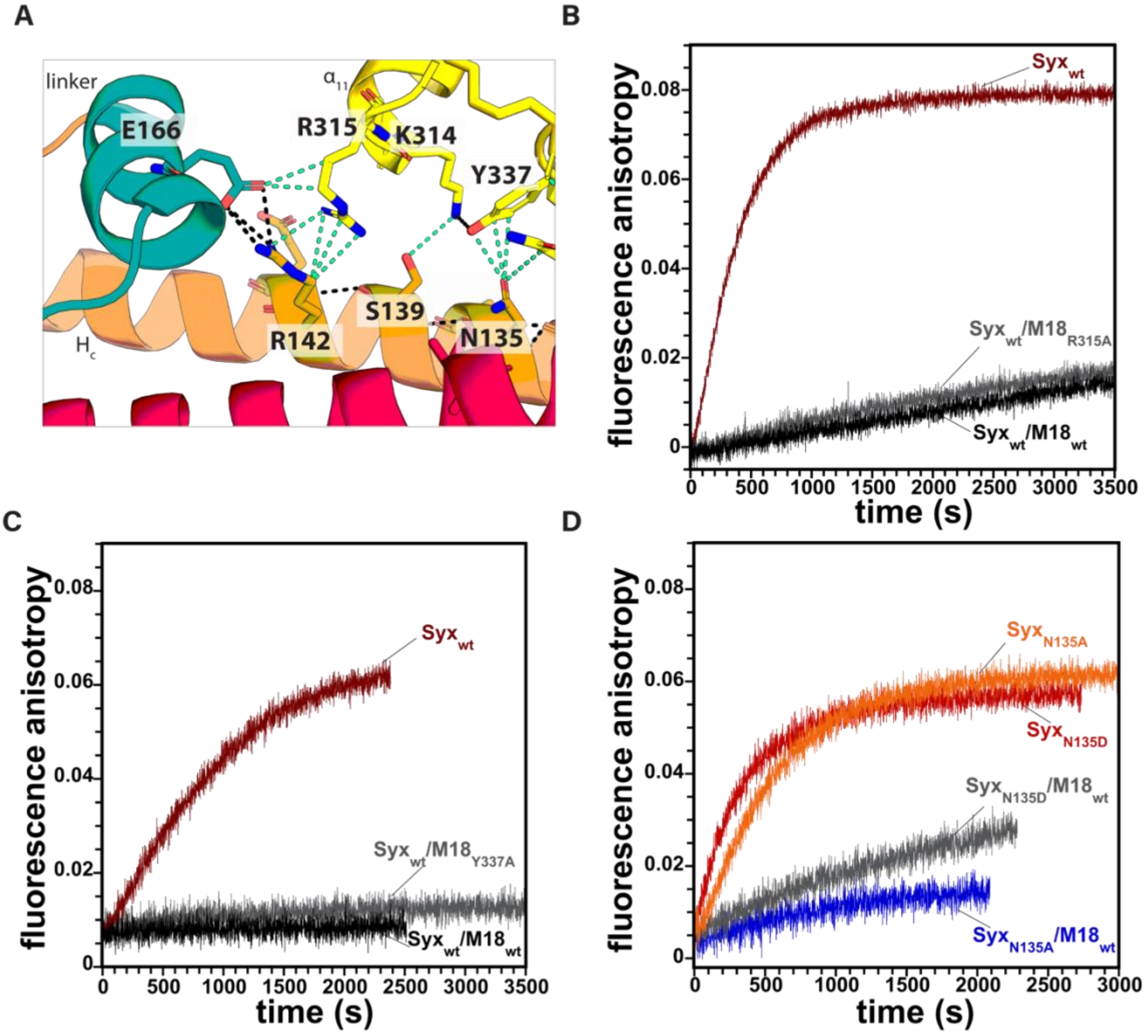


**Figure S2: The inhibitory effect of Munc18-1 on SNARE assembly is affected by mutations at the tip of Domain 3a of Munc18-1.**

**(A)** Protein interface interactions at the Munc18-1 Domain 3a and syntaxin-1a linker region. Black dashed lines indicate polar contacts and cyan dashed lines indicate any other interaction within 4.0 Å. In the Munc18-1/syntaxin-1a complex (pdb: 3c98), the tip of Domain 3a interacts with the Hc–linker–H3 regions of syntaxin-1a (1,2). R315 of Munc18-1 is in polar contacts with E166 and R142 of syntaxin-1a. Y337 of Munc18-1 is also involved in the electrostatic interactions with N135 of syntaxin-1a's Hc helix. **(B)** A mutation of R315 to alanine, $M18_{R315A}$, did not significantly reduce the inhibition of SNARE complex assembly of the bound syntaxin-1a (grey). **(C)** Mutation of Y337 to alanine, $M18_{Y337A}$, also did not reduce the inhibition of SNARE complex assembly of the bound syntaxin-1a significantly. **(D)** Mutation of N135 to an alanine, $Syx_{N135A}$, did not significantly reduce the inhibition of SNARE complex assembly by Munc18-1 significantly. Inhibition was reduced when N135 was substituted by an aspartate ($Syx_{N135D}$; grey trace). Mixing experiments were carried out as described in the legend of Fig. 2.

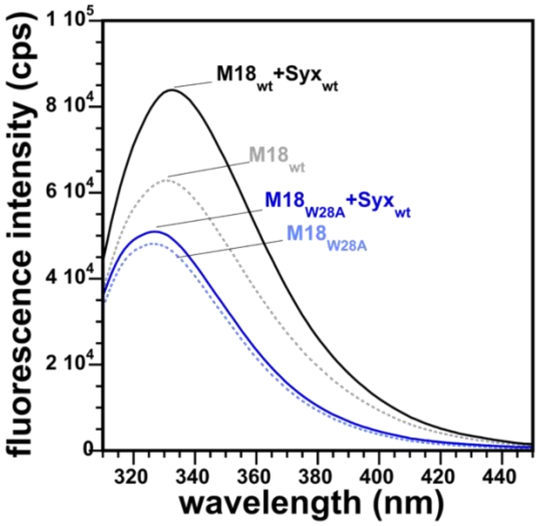


**Figure S3: The increase in intrinsic tryptophan fluorescence upon the formation of the Munc18-1/syntaxin-1a complex originates from tryptophan 28 of Munc18-1.**

Fluorescence emission spectra of individual $M18_{wt}$ and $M18_{W28A}$ (200 nM) before (grey dotted and light blue dotted line) and after addition of an excess of $Syx_{wt}$ (300 nM) (black and blue, respectively) were recorded upon excitation at 295 nm. As expected, the intrinsic fluorescence of $M18_{W28A}$ reduced compared with $M18_{wt}$ . Upon addition of $Syx_{wt}$ to $M18_{wt}$, an increase in tryptophan fluorescence was noted. Only a small change was observed when $Syx_{wt}$ was added to $M18_{W28A}$. Note that Munc18-1 possesses five tryptophans, while syntaxin-1a has no tryptophan.

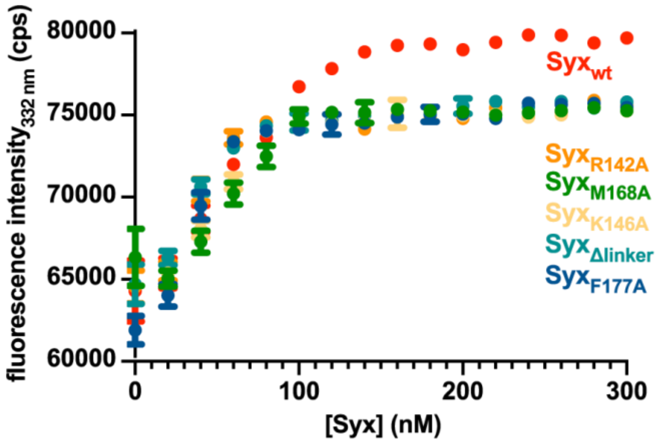


**Figure S4: Changes in tryptophan fluorescence upon titration of different syntaxin-1a variants to Munc18-1.**
The addition of all syntaxin-1a variants to Munc18-1 (150 nM) led to a clear increase in fluorescence that was saturated at an equimolar ratio. A higher fluorescence maximum was reached for Syx$_{wt}$ compared with the linker variants. Fluorescence was measured at 332 nm upon excitation at 295 nm.

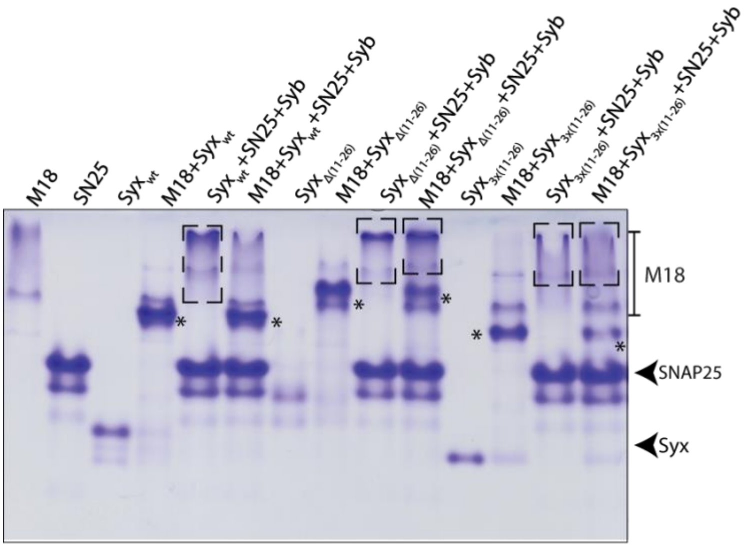


**Figure S5: Munc18-1$_{wt}$ forms stable complexes with syntaxin-1a N-terminus variants.**

In total, 55 µmol of Munc18$_{wt}$ and 34 µmol of Syx$_{wt}$, Syx$_{Δ(11-26)}$, and Syx$_{3x(11-26)}$ were incubated for 30 min at room temperature in the presence or absence of the SNARE proteins SNAP25 (SN25) and synaptobrevin (Syb) at 90 µmol each. Complex formation was monitored by native gel electrophoresis. The positions of syntaxin and SNAP25 are indicated by arrows. Dashed boxes indicate the broad band corresponding to the SNARE complex, and asterisks indicate the M18/Syx complexes. All syntaxin-1a variants were able to form SNARE complexes with SNAP-25 and synaptobrevin and complexes with Munc18-1. Munc18-1 did not completely block the formation of the SNARE complex for the *N*-terminus mutants, as complex formation was observed in the presence of Munc18-1. Note that a smear corresponding to SNARE complex is observed also for Syx$_{wt}$ in the presence of M18; however, this is probably an excess of uncomplexed Syx$_{w}$, as seen by the residual free Syx$_{wt}$ also present the lane of M18+Syx$_{wt}$. M18/Syx$_{3x(11-26)}$ was the least stable in the presence of SNARE interactors, as the band corresponding to the complex was fainter in the presence of SNAP-25 and synaptobrevin.

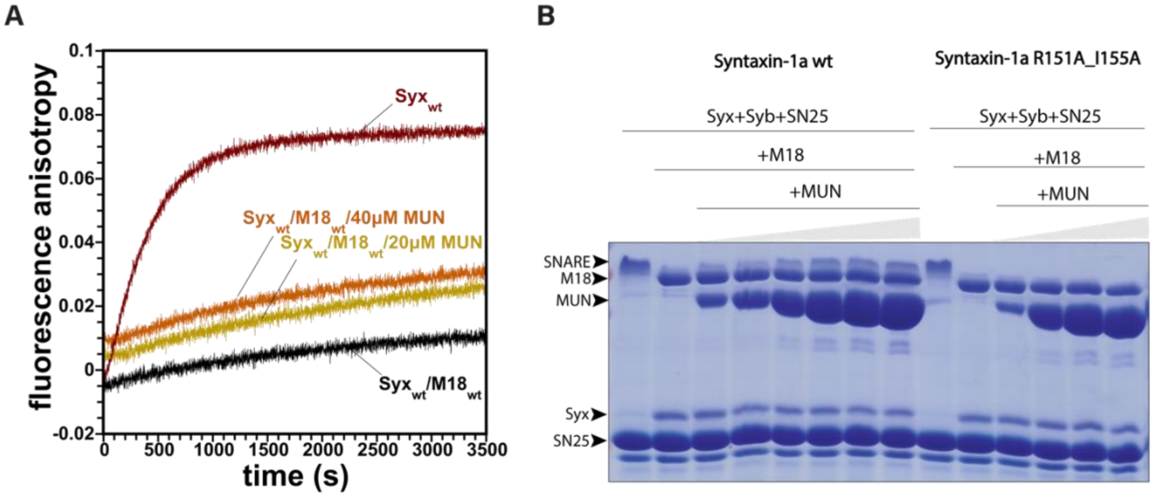


**Figure S6: SNARE complex inhibition by Munc18-1 in the presence of the MUN domain.**

**(A)** Fluorescence anisotropy of premixed $M18_{wt}$, $Syx_{wt}$, and Syb*[28] mixtures in the absence (black) and the presence of the MUN domain (20 µM (ochre) and 40 µM (orange)). Reactions were started by adding SNAP-25. No significant change in the speed of SNARE complex assembly was noted, although a higher fluorescence anisotropy starting value was observable in the presence of the MUN domain. This could be the result of crowding caused by the high concentration of the MUN domain used. Mixing experiments were carried out as described in the legend of Fig. 2. **(B)** $Syx_{wt}$ or $Syx_{R151A\_I155A}$ (16 µmol) was mixed with $M18_{wt}$ (72.8 µmol), SNAP25 (82 µmol) and Syb (82 µmol), and increasing amounts (9, 16, 41, 65, 82, and 98 µmol) of the MUN domain of Munc13-1. The mixtures were then subjected to SDS-PAGE without boiling of the sample. After the run, the gel was stained with Coomassie Blue. Note that the bands corresponding to the SDS-resistant complex ran higher than that of Munc18-1 but overlap somewhat. Increasing amounts of the MUN domain appeared to lead to a stronger band of the SDS-resistant SNARE complex. Increased formation of the SNARE complex was not observed for the syntaxin variant $Syx_{R151A\_I155A}$.

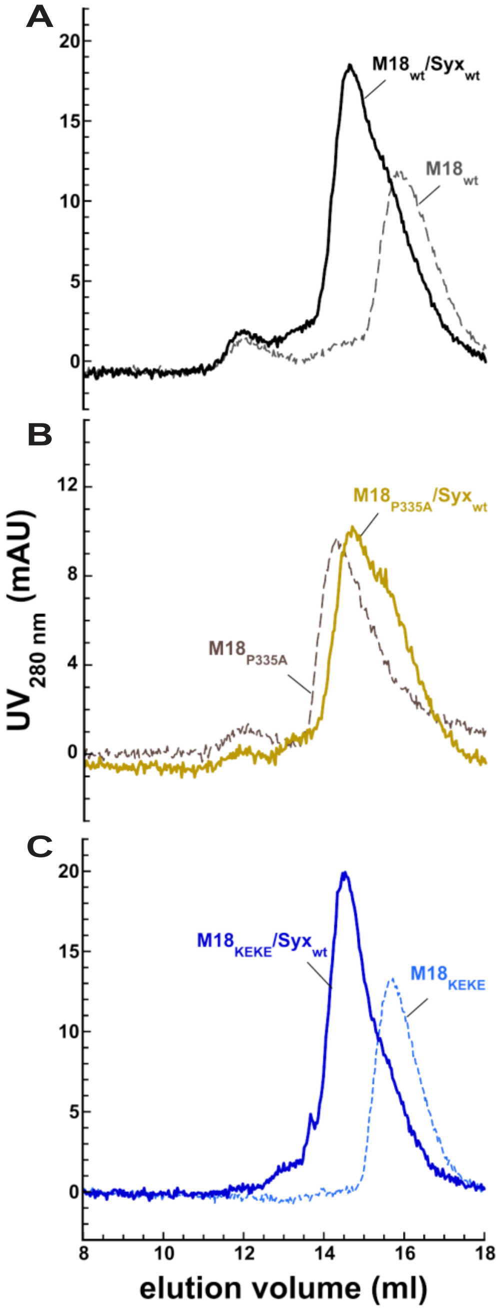


**Figure S7: Analysis of complex formation between different Munc18-1 variants and syntaxin 1a by size exclusion chromatography.**
Prior to separation on a Superdex 200 column, the proteins (≈22 µM each) were incubated for 30 min at room temperature. Protein elution was monitored at 280 nm. Elution profiles of **(A)** $M18_{wt}$ alone (grey dashes) and mixed with $Syx_{wt}$ (black), **(B)** $M18_{P335A}$ alone (brown dashes) and mixed with $Syx_{wt}$ (ochre), and **(C)** $M18_{P335A}$ alone (blue dashes) and mixed with $Syx_{wt}$ (blue). All Munc18-1 variants comigrated with syntaxin-1a. Note that individual $M18_{P335A}$ eluted earlier than other Munc18-1 variants, suggesting that it formed an oligomer, most probably a dimer.

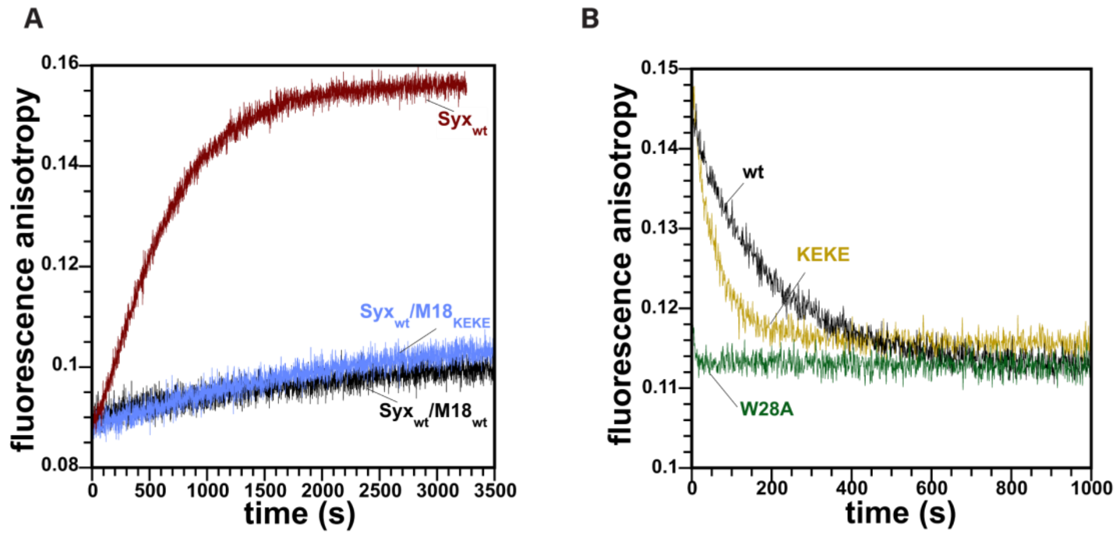


**Figure S8: Like Munc18-1$_{wt}$, Munc18-1$_{KEKE}$ also inhibits the formation of the SNARE complex.**

**(A)** SNARE complex formation is inhibited in the presence of M18$_{KEKE}$ (violet trace) or M18$_{wt}$ (black trace). Formation of the SNARE complex was monitored by fluorescence anisotropy of Syb$^{*28}$. In the absence of Munc18, SNARE complex formation can be seen by the increase in fluorescence anisotropy (brown trace). Mixing experiments were carried out as described in the legend of Fig. 2. **(B)** The M18$_{KEKE}$/Syx$_{wt}$ complex dissociated faster than the M18$_{wt}$/Syx$_{wt}$ complex. Competitive dissociation experiments were carried out as described in the legend of Fig. 5D. An excess of unlabeled Syx1a (5 µM) was added to a premix of Munc18 with Oregon Green-labeled Syx1a. Interestingly, the dissociation of the M18$_{W28A}$/Syx$_{wt}$ complex was the fastest among all the complexes studied. In fact, it was difficult to determine the exact value for this reaction, as the anisotropy decayed too fast for the hand-mixing approach used. The dissociation rates are given in Table S1.

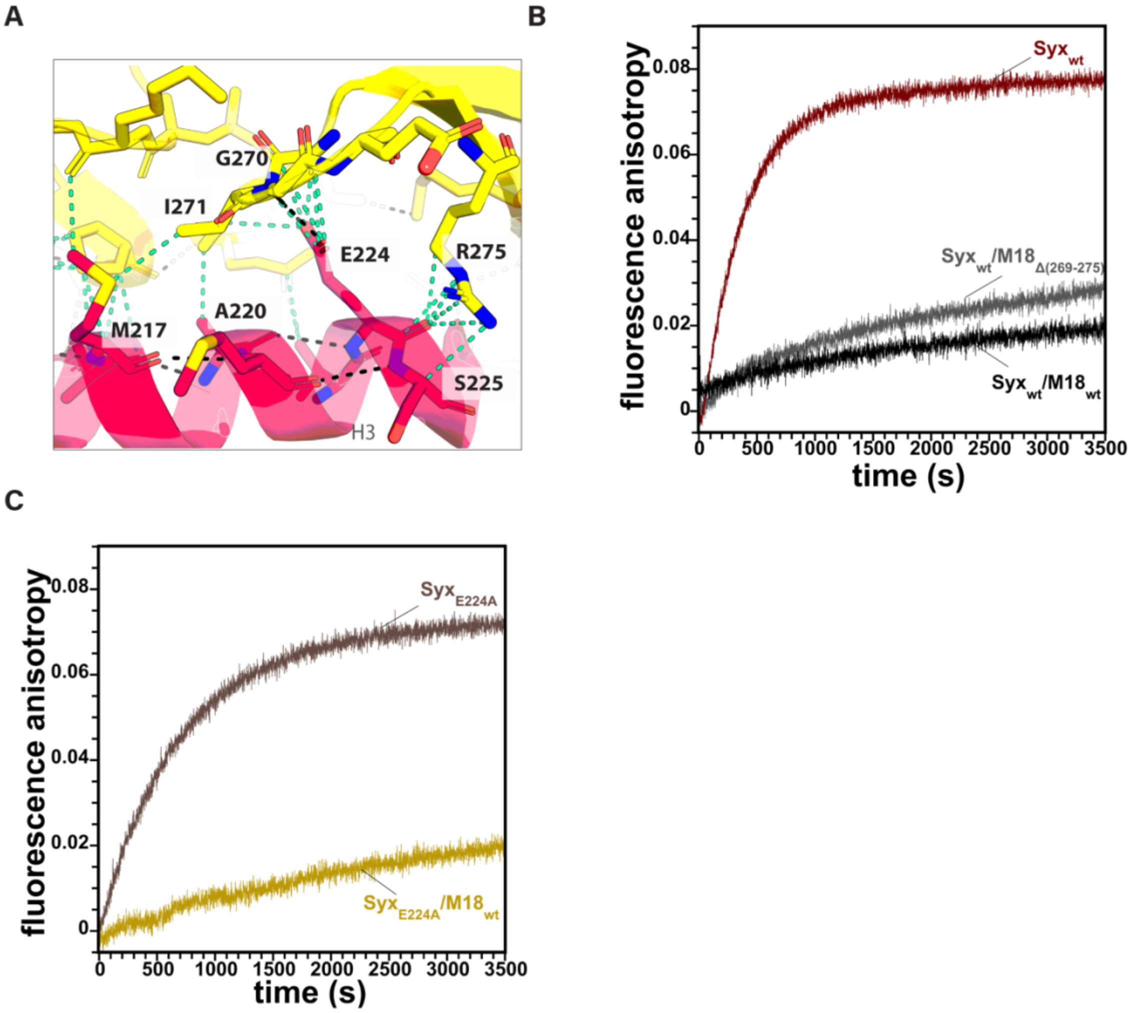


**Figure S9: Munc18-1 $\beta_{10}\beta_{11}$ loop residues contribute to Munc18-1's inhibition of the formation of the SNARE complex exerted by Munc18.**

**(A)** Enlarged section of the Munc18-1/syntaxin-1a complex structure, highlighting the interface of the $\beta_{10}\beta_{11}$ loop (aa 269–274) with the residues M216, A220, E224, and S225 of syntaxin-1a H3. The interacting residues are shown as sticks and are labeled. Black dashed lines indicate polar contacts and cyan dashed lines indicate any other interaction within 4.0 Å. **(B)** The SNARE complex forms somewhat faster in the presence of $M18_{\Delta 269\text{-}275}$ (grey trace) than in the presence of $M18_{wt}$ (black trace). **(C)** Munc18 inhibits the formation of the SNARE complex of $Syx_{E224A}$ (ochre trace), as monitored by fluorescence anisotropy.

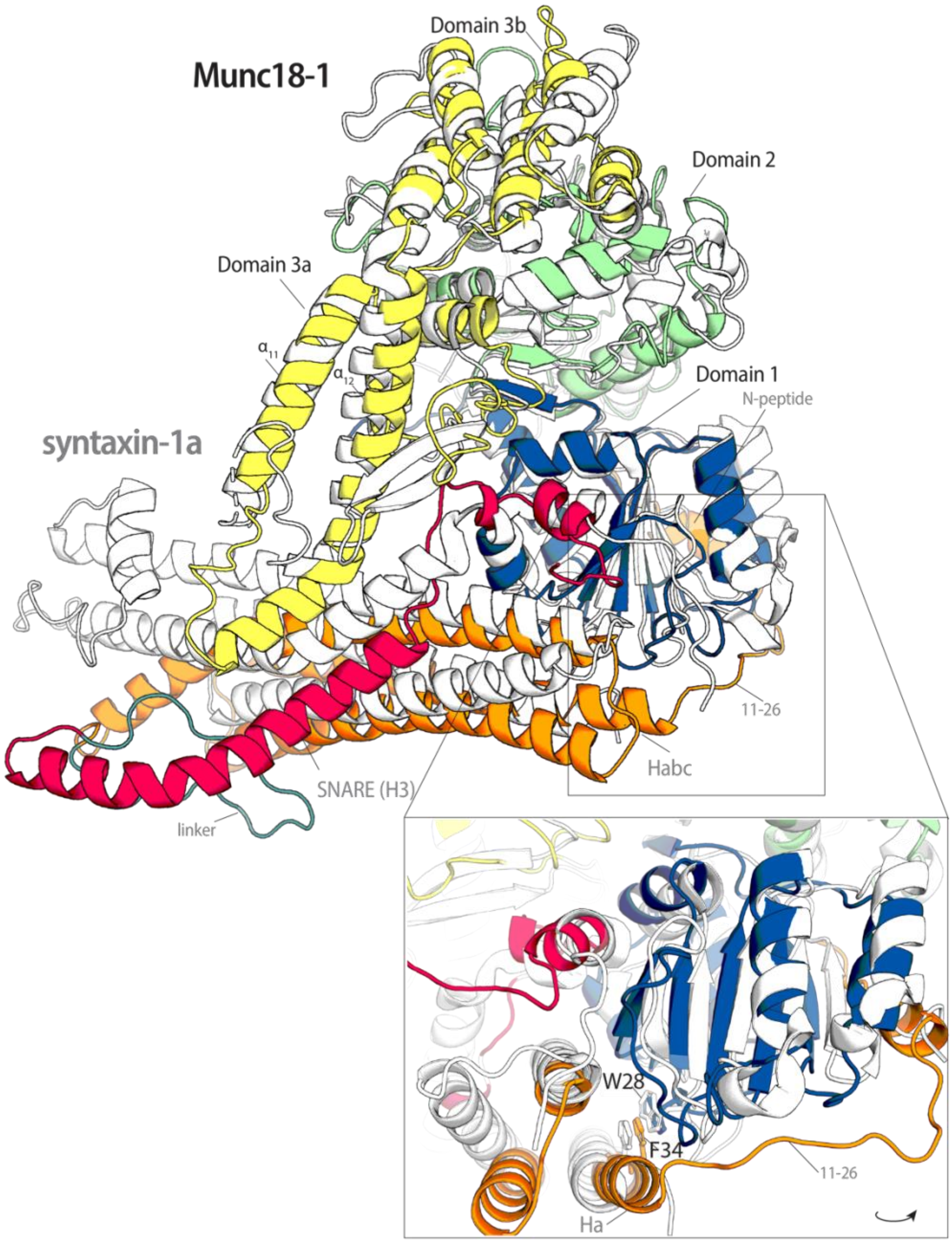


**Figure S10: Visualization of the putative conformational change in the Munc18-1/syntaxin-1a complex.**

**Top:** Overlay of the modeled structure of the Munc18-1/syntaxin-1a complex in a more open conformation (color code as in Fig. 6) and the crystal structure (in white). **Bottom:** Close-up side view of Domain 1 of Munc18-1, bound on two sides by the Habc domain and the N-peptide of syntaxin-1a. The positions of W28 and F34 are indicated. Note that the linker region, aa 11–26, was modeled but was not resolved in the crystal structure.

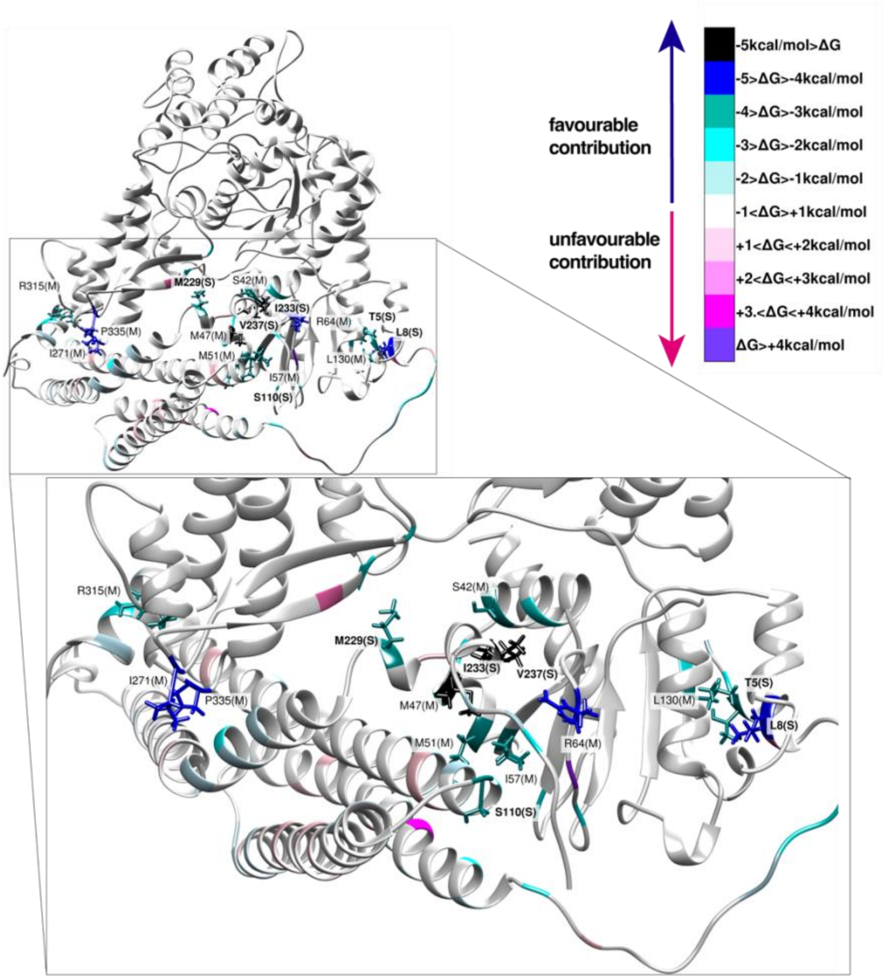


**Figure S11: Contribution of residues to the free energy of the interaction (ΔG) between Munc18-1 and syntaxin-1a.**

The free energy was determined by the MM-GBSA method (3). Residues are shaded from black (most favorable contribution) to white (negligible contribution) to violet (most unfavorable contribution). The residues with the most favorable contributions are represented as stick and have been labeled. The syntaxin-1a residues are labelled in bold, whereas the Munc18-1 residues are labeled in normal font.

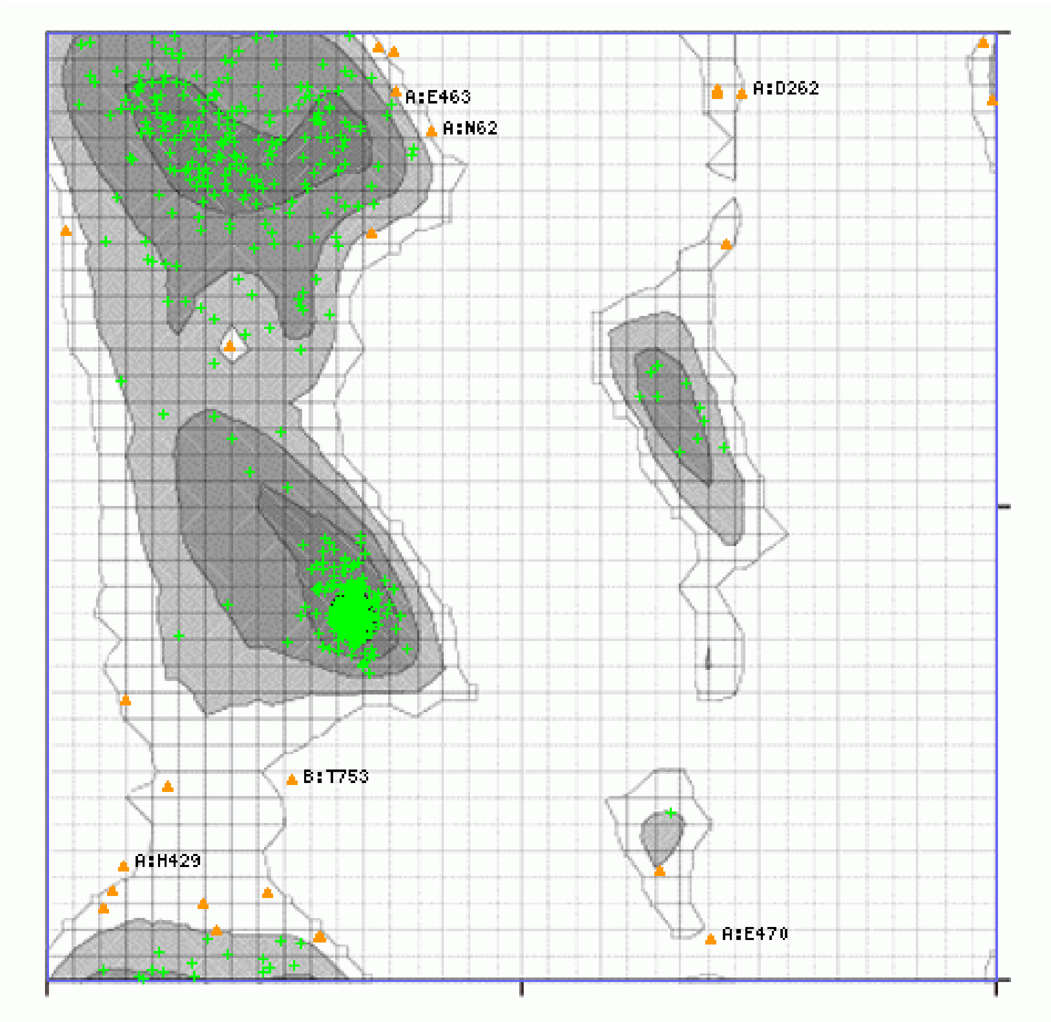


**Figure S12: Ramachandran plot of the modelled Munc18-1/syntaxin-1a complex in intermediate conformation.**
The torsional angles of the amino acids φ (phi) and ψ (psi) for the model of the Munc18-1/syntaxin-1a complex are shown according to (4). In total, 750 residues (96.65 %) were found within favourable regions (green crosses), and 26 residues (3.35 %) belong to the preferred regions (orange triangles). The torsional angles of the 31 proline and 30 glycine residues are not shown.

**Table S1. Competitive dissociation rates of Munc18-1/syntaxin-1a complexes.**
Note that it was impossible to determine the exact value for the $M18_{W28A}/Syx_{wt}$ off-rate, as the anisotropy decayed too fast for the hand-mixing approach used. We estimate it to be faster than 0.1/s.

| Munc18-1 variant | $K_{off}$ |
|---|---|
| wt | 0.004/s |
| K332E/K333E | 0.014/s |
| Δ269-275 | 0.025/s |
| Y337A | 0.039/s |
| Δ317-333 | 0.045/s |
| P335A | 0.061/s |
| W28A | n.d. |

**Table S2. Contributions of the residues to the free energy of interaction between Munc18-1 and syntaxin-1a.**
The energy contributions were estimated by the MM-GBSA method based on 100 conformations, and the stability was estimated by FoldX 5.0 (5,6).

| Molecule | Residue | Average contribution to interaction (kcal/mol) | Contribution to stability (kcal/mol) |
|---|---|---|---|
| syntaxin-1a | F34 | -2.9 | -1.67 |
| | N135 | unfavourable 0.10 | unfavourable 0.59 |
| | R142 | unfavourable 0.71 | -0.01 |
| | K146 | unfavourable 1.25 | unfavourable 1.48 |
| | E163 | -0.2 | unfavourable 3.34 |
| | E164 | -1.34 | unfavourable 1.38 |
| | L165 | -0.11 | -1.56 |
| | E166 | -0.96 | unfavourable 1.41 |
| | D167 | -2.88 | unfavourable 2.75 |
| | M168 | -0.07 | -0.48 |
| | L169 | -0.12 | -2.00 |
| | E170 | -1.68 | unfavourable 1.46 |
| | F177 | unfavourable 0.003 | -3.59 |
| Munc18-1 | W28 | -2.95 | -0.41 |
| | W288 | -0.04 | -3.79 |
| | R315 | -3.47 | unfavourable 0.29 |
| | K332 | -0.01 | unfavourable 0.07 |
| | K333 | -0.43 | -0.11 |
| | P335 | -4.22 | -1.49 |
| | Y337 | -0.07 | -1.15 |
